# Modeling the impact of single-cell stochasticity and size control on the population growth rate in asymmetrically dividing cells

**DOI:** 10.1101/2020.11.30.404012

**Authors:** F. Barber, J. Min, A. W. Murray, A. Amir

## Abstract

Microbial populations show striking diversity in cell growth morphology and lifecycle [1]; however, our understanding of how these factors influence the growth rate of cell populations remains limited. We use theory and simulations to predict the impact of asymmetric cell division, cell size regulation and single-cell stochasticity on the population growth rate. Our model predicts that coarse-grained noise in the single-cell growth rate λ decreases the population growth rate, as previously seen for symmetrically dividing cells [2]. However, for a given noise in λ we find that dividing asymmetrically can enhance the population growth rate for cells with strong size control (between a “sizer” and an “adder”). To reconcile this finding with the abundance of symmetrically dividing organisms in nature, we propose that additional constraints on cell growth and division must be present which are not included in our model, and we explore the effects of selected extensions thereof. Further, we find that within our model, epigenetically inherited generation times may arise due to size control in asymmetrically dividing cells, providing a possible explanation for recent experimental observations in budding yeast [3]. Taken together, our findings provide insight into the complex effects generated by non-canonical growth morphologies.

## 1 Introduction

Recent years have expanded our understanding of heterogeneity at the single cell level, with clonal populations displaying variability in a range of physiological parameters, including cell generation times (the time between cell birth and division), cell size and gene expression [4, 5, 6, 7, 8]. This revolution in single-cell microbiology drove a renewed interest in the effect of heterogeneity on cell fitness, taken here to be described by the population growth rate [2, 9, 10, 11]. In contrast, a relatively unexplored factor affecting cell fitness is cell growth morphology; microbial cells display an astonishing degree of variability in growth morphology and life cycle, ranging from symmetric division in the vegetative growth of bacteria such as *Bacillus subtilis* and *Escherichia coli*, to the asymmetrically dividing, budding yeast *Saccharomyces cerevisiae*, to the diverse growth morphologies observed recently in a range of marine yeasts [1]. However, our understanding of the physiological effect of division asymmetry on the population growth rate remains limited.

Early work demonstrated that the population growth rate Λ_p_ obeys the Euler-Lotka equation [12]

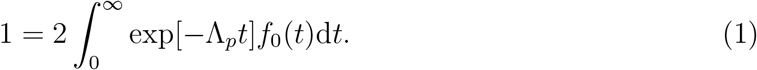

Here *f*_0_ (*t*) is the distribution of generation times measured by tracking all cells in a growing population (called the lineage tree or tree distribution here), illustrated in Figure 1 (A) [13, 2]. If generation times are uncorrelated between related cells (the independent generation time or IGT case), *f*_0_(*t*) is the same as the distribution obtained from tracking cells along a single cell lineage, however, correlated generation times have been observed in a range of organisms [7, 8, 3, 14, 15], meaning the full tree distribution is required for Equation 1 to hold. These generation time correlations are expected as a direct consequence of size control, whereby cells couple their growth and division to constrain the spread of sizes observed throughout a population [16]. Including size control when modeling cell cycle progression fundamentally changes the predicted impact of stochasticity at the single-cell level on Λ_*P*_ [2]. Prior studies that did not incorporate cell size control have concluded that noise in generation times can enhance the population growth rate [3, 9]. In contrast, studies incorporating cell size control predict that the single cell exponential growth rate λ sets Λ_*P*_, with Λ_*P*_ = λ exactly in the absence of noise in their single cell growth rate [2]. This can be readily shown by requiring that the cell size distribution reaches steady state with a constant average size, since 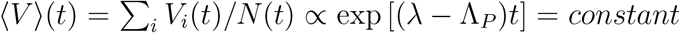. Coarse-grained noise in λ then decreases ΛP below the average single cell growth rate, while for a given noise in *λ*, increasing noise in generation times is predicted to only have a smaller, secondary effect [2, 10].

**Figure 1:**
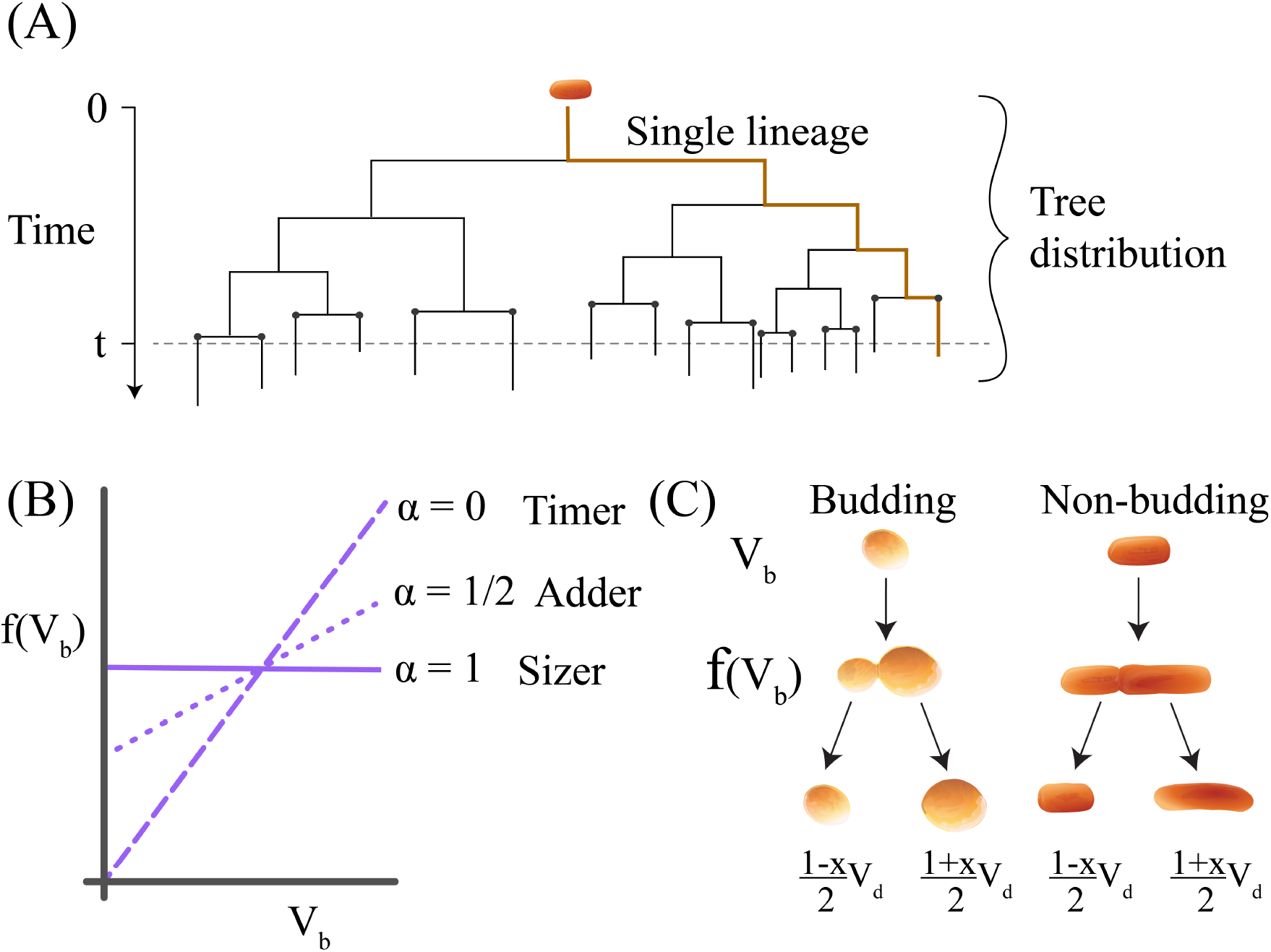
(A) Illustration of the tree distribution for a growing population of cells, ter minated at a time-point t but including those cell cycles that are unfinished at *t*. A single lineage is shown in orange. Each node corresponds to a single cell division event. Grey dots correspond to the final division events before *t*. (B) Illustration of differences in cell size control policy. (C) Illustration of asymmetric division in different growth morphologies. Budding cells set the plane of division early on in the cell cycle and direct growth to a newly forming bud beyond that division plane. In contrast, non-budding cells set the plane of division when division occurs, meaning that growth throughout the cell cycle affects both progeny.

Asymmetric cell division generates two distinct cell types; in budding yeast these are known as daughters (the smaller cells) and mothers (the larger cells). To compensate for this difference in size, daughters have a longer average generation time than mothers. One early study in the context of budding yeast obtained a theoretical prediction for ΛP under the assumption of constant division times for daughters (*τ_D_*) and for mothers (*τ_M_*), with *τ_M_* ≤ *τ_D_* [14] (Section S3). A more recent study considered the effect of correlated generation time noise on the population growth rate of budding yeast cells [3]. However, as discussed above, this work did not employ a model of cell size control, leading the authors to predict that single cell stochasticity and epigenetically inherited generation times can enhance the population growth rate. Our results disagree with these predictions. Here we show that for cells that regulate their size, the population growth rate is set primarily by the single cell growth rate, with noise in the single cell growth rate *decreasing* the population growth rate, as in the case of symmetrically dividing cells. We further show that asymmetric division can *increase* the population growth rate, and that epigenetically inherited generation times can arise as a natural consequence of size control in asymmetrically dividing cells.

## 2 Results

### 2.1 Model for asymmetric population growth

As discussed above, when the single cell growth rate λ is constant, Λ_*P*_ = λ exactly [2]. To study the effect of finite noise in λ, we modeled the growth of two coupled cell populations (*N_M_* for mothers and *N_D_* for daughters). The growth of these populations is described in the limit of large population numbers by

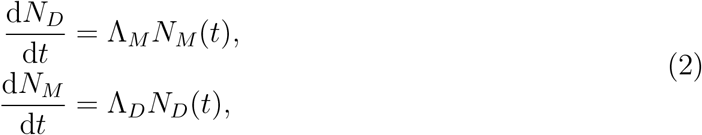

since a cell of either type divides to give one new cell of each type. Here Λ_*D*_ and Λ_*M*_ correspond to the division rates per cell of cell types *D* and *M* respectively. Assuming steady state composition of the population, with a constant relative difference in the number of different cell types *m*(*t*) ≡ (*N_D_* (*t*) — *N_M_* (*t*))/(*N_D_* (*t*) + *N_M_*(*t*)) = *m* (which we will corroborate later), the full population *N*(*t*) = *N_D_*(*t*) + *N_M_*(*t*) will grow exponentially with growth rate 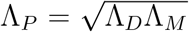 (Section S1). Importantly, the Euler-Lotka equation for the two population system described above still holds (Section S2):

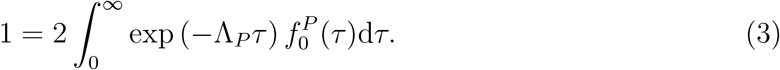

Here 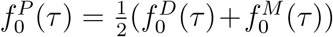 is the distribution of generation times measured over the full lineage tree, including both mother and daughter cells. A corresponding constraint equation also exists for the relative difference in population numbers *m* (Equation S12). We note that although *m* will in general be greater than zero, with a larger fraction of daughter cells than mother cells at a given point in time, the populations of daughter and mother cells will be equal when measured over the full lineage tree, leading to the factor of 1/2 in the definition of 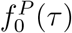. Equation 3 can also be shown to apply in the case of finite population sizes using the transport equation approach outlined in Ref. [17]. Our current approach provides the additional benefit of predicting the ratio of cell types m present in the population at a single time-point, given the distribution of interdivision times.

### 2.2 Models of size control

To study the effect of size control on Λ_*P*_, we define a growth function *h*(*V_b_*) that sets the target volume at division to be a linear function of volume at birth *V_b_*, with a tunable parameter *α* [18]:

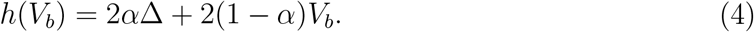

Setting *α* = 0 gives a timer model, in which cells grow to double their volume between birth and division, while *α* =1/2 gives an adder model with a constant volume Δ added between birth and division, and *α* =1 gives a sizer model where cells grow to a threshold size 2Δ at division (see Figure 1 (B)). Cell volume at division is then given by *V_d_* = *h*(*V_b_*) exp [λ*η*], with associated generation time *t* = ln |*h*(*V_b_*)/*V_b_*|/λ + *η* where 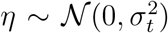 is a coarse grained noise in generation times (independent and identically distributed, I.I.D., for each newborn cell) and λ is the I.I.D. exponential single cell growth rate taken to be 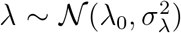. We define the parameter *x* as the relative difference in volume at birth between the daughter and mother cells produced from a given division event: 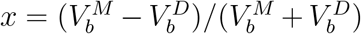 [19], as described in Figure 1 (C). This implies 0 < *x* < 1. We will use subscripts *b* and *d* to denote whether the cell volume is evaluated at birth or at division, while the superscripts *D* and *M* correspond to the two different cell types. When a statement is independent of cell type we use the superscript *P* to denote that cell. Our prior work has studied the differences between budded cells and non-budded cells as shown in Figure 1 (B) [20]. We compared simulations of budding vs. non-budding cells, finding no observable differences for the population growth rate in cases without generation time noise, and only minor differences in cases with nonzero generation time noise (see Figure S1 (A)). Since our analytical model describes non-budding cells, our results are presented for non-budding cells unless stated otherwise. We now study the effects of variation in division asymmetry *x*, size control strategy *α* and noise terms *σ_λ_* and *σ_t_* on *Λ_P_*.

### 2.3 An approximate solution for the population growth rate

Solving Equation 3 to infer the population growth rate for a general size regulation model is difficult due to correlations between successive generation times. These correlations vanish for symmetrically dividing cells without noise in generation times that follow any mode of size control, as has been studied previously [2]. These correlations also vanish for asymmetrically dividing cells following a sizer model (*α* =1) without noise in generation times, allowing us to obtain an approximate solution in this case:

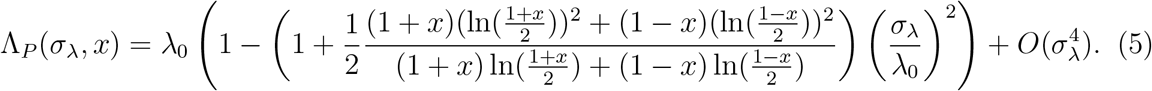

See Section S3 for details. Setting *x* = 0 recovers the approximate solution for symmetric growth [2], with

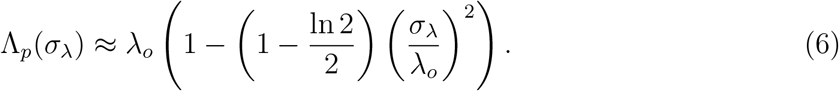

From Equation 5, we predict that noise in *λ* will tend to decrease the population growth rate as in the case of symmetric division [2]. However, our model further predicts that for non-zero *σ_λ_*, Λ_*P*_ can be increased by increasing the division asymmetry. We tested these predictions by simulating the growth of populations of cells, using Equation 4 to regulate the timing of division (see Methods for details). Figure 2 (A) demonstrates the independence of Λ_*P*_ from *σ_t_* or *x* for *σ_λ_* = 0. The figure shows the case of adder cells (*α* = 0.5), but this result is the same for any strategy of size control. Figure 2 (A) also shows that finite *σ_λ_* decreases the population growth rate further below 〈λ〉, while noise in generation times *σ_t_* has a small, secondary effect in the range of biologically relevant division asymmetry values. The negative impact of noise in the single cell growth rate on the population growth rate is on the order of 1.5% for the biologically relevant case of *σ_λ_/λ*_0_ ≈ 0.15 [7], indicating that this effect may be significant from an evolutionary standpoint. We note that the effect of *σ_t_* becomes more substantial in the regime of extreme division asymmetry, as shown in Figure S1 (A), however, this regime is not believed to be biologically relevant based on experimental measurements of the division asymmetry x in budding yeast ranging from 0.2 to 0.35 across different growth conditions [8]. We tested Equation 5 for a sizer model against simulations across a range of values for *σ_λ_* and *x* (with *σ_t_* = 0), finding consistently good agreement as shown in Figure S1 (B). These findings make the strong prediction that increasing division asymmetry can enhance Λ_*P*_ for cells that regulate their size with a sizer strategy and have non-vanishing growth rate variability.

**Figure 2:**
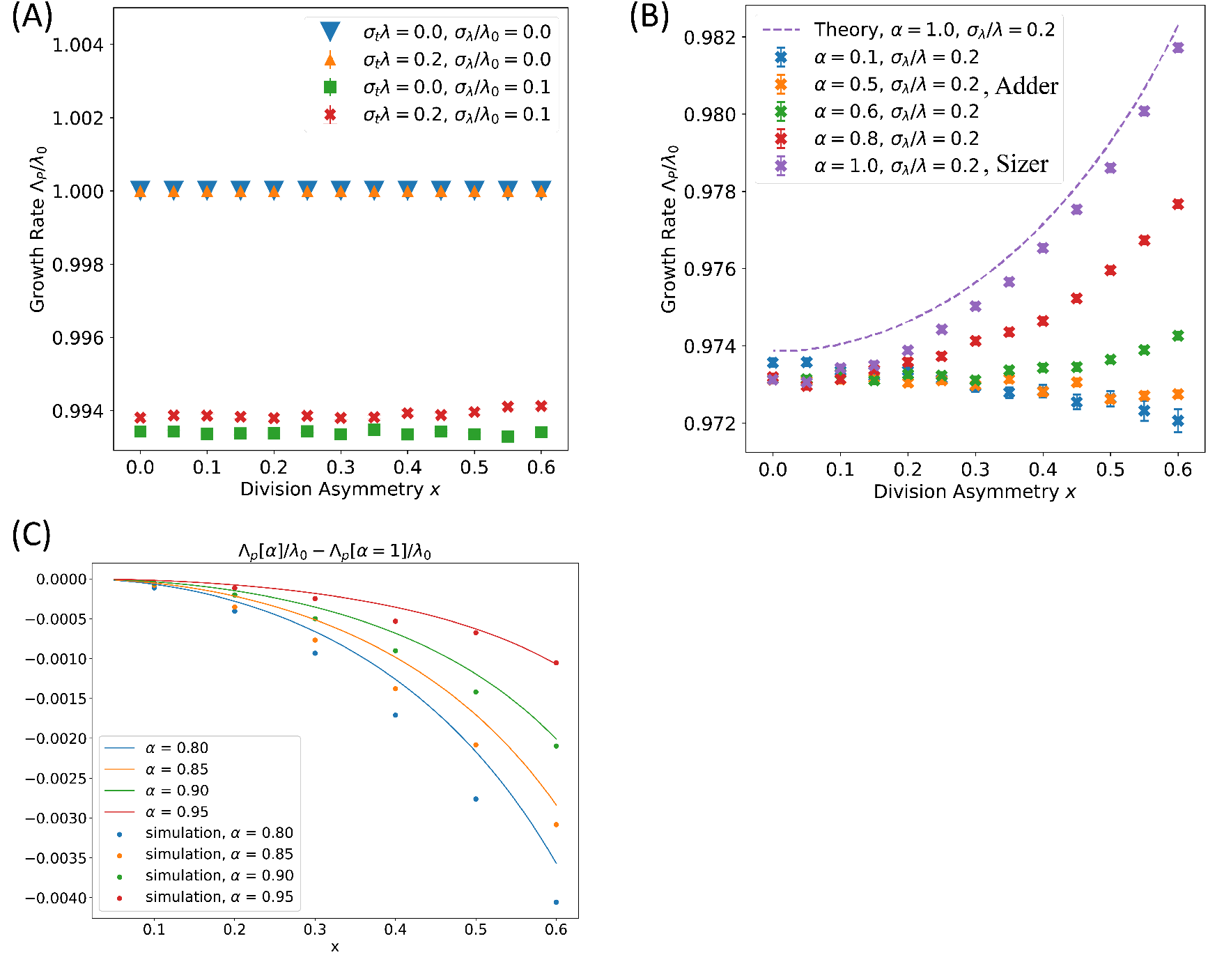
The population growth rate Λ_*P*_ is dependent on noise in single cell growth rate *σ_λ_*, the division asymmetry *x* and the size control strategy *α* for asymmetrically dividing cells. (A) Λ_*P*_ = λ exactly in the absence of noise in λ for cells that display size control, regardless of time-additive noise in generation times or division asymmetry. Coarse-grained noise in λ decreases Λ_*P*_. Plot is shown for adder cells with *α* = 1/2. (B) Λ_*P*_ plotted for a range of size control strategies *α* with *σ_λ_*. Size control strategies weaker than an adder do not display any benefit in Λ_*P*_ from dividing asymmetrically. *σ_λ_*/*λ*_0_ = 0:2, *σ_t_* = 0. (C) Deviation from a sizer causes a relative decrease in Λ_*P*_ for large *x*. The difference between Equation S43 for Λ_*P*_ (*α*) and Equation 5 for Λ_*P*_ (*α* = 1) is plotted against *x* for deviations from *α* = 1. Parameters are listed in the Figure legend. Data points correspond to simulations, while lines represent theoretical predictions. Error bars show the standard error of the mean.

To explore whether increasing division asymmetry consistently increased Λ_*P*_ for different strategies of cell size control within our model, we simulated population growth across a range of α values between 0 and 1. Results are plotted in Figure 2 (B), showing that for cells that divide asymmetrically, the growth rate gain associated with increasing x shown in Figure 2 (B) is reduced for size control strategies weaker than a sizer, while for *α* ≤ 1/2 (size control weaker than an adder model), increasing x has a slight tendency to *decrease* Λ_*P*_. Cells following an adder strategy showed little dependence of Λ_*P*_ on division asymmetry. This strong dependence of Λ_*P*_ on the strategy of size control has not been observed previously in studies focusing on symmetric division, and to our knowledge is the first instance in which Λ_*P*_ depends on the strategy of size control in exponentially growing cells.

By expanding around *α* = 1 we obtained an approximate expression for the growth rate Λ_*P*_(*α*) for small |*α* — 1| (see Section S4 and Equation S43 for details). Figure 2 (C) shows our predictions for the difference between Λ_*P*_(*α*) and Λ_*P*_ of a sizer, We observe good agreement with simulations for small *x*, supporting our result that asymmetrically dividing cells with *α* < 1 will have a lower Λ_*P*_ relative to the *α* =1 sizer case.

For completeness we also explored the behavior of the population asymmetry factor *m*, showing that in the case of a sizer model *m = x* exactly, independent of *σ_λ_* (Section S3 and Figure S2), and that weaker strategies of size control show a weaker dependence of *m* on *x*.

### 2.4 Generation time correlations

One recent study observed positively correlated generation times in closely related budding yeast cells [3]. When these correlations were introduced in simulations of growing populations of cells that did not regulate their size, they led to an enhancement of the population growth rate Λ_*P*_. This prompted the authors to conclude that the epigenetic inheritance of generation times may enhance the population growth rate. The experimental observation of positively correlated generation times is surprising when contrasted with the negative correlations associated with cell size control in symmetrically dividing cells [2]. To investigate this, we adopted a model of cell cycle duration (Equation 7) that has been previously applied to analytically calculate the generation time correlation coefficients of cells growing with varying strategies of size control [16]:

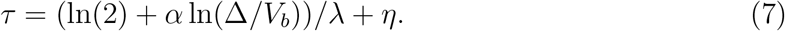

Here Δ is the mean cell size at birth, 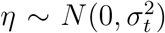 is a coarse grained I.I.D. noise in generation times [16], and 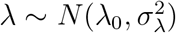 is the noisy single cell growth rate. Equation 7 arises from the growth policy 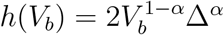, which agrees to first order with Equation 4 when Taylor expanded around the average newborn size Δ. *α* = 0 corresponds to a timer and *α* =1 corresponds to a sizer, and *α* = 0.5 corresponds to first order with an adder. Using this model we obtained an approximate formula for the Pearson correlation coefficient (PCC) arising from cell size control in asymmetrically dividing cells in the case without noise in the single cell growth rate (*σ_λ_* = 0) (see Section S8 for details), finding that positively correlated generation times can arise as a natural consequence of cell size control. This counter-intuitive result for asymmetrically dividing cells contrasts strongly with the negative generation time correlation *PCC* = — *α*/2 that is predicted by this model for symmetrically dividing cells, but this discrepancy can be readily explained. Negatively correlated generation times arise when symmetrically dividing cells time their division events to correct for noise-induced fluctuations in cell size that were generated in previous cell cycles. For asymmetrically dividing cells there is an additional, positive term that arises due to a cell’s lineage: a daughter cell that is born from a daughter cell will be smaller than the average daughter cell. In the case of an adder model, this smaller daughter cell will take a longer time to add the same volume increment Δ through exponential growth, leading to a longer than average division time. Conversely, a daughter cell that is born from a mother cell will be larger than an average daughter cell, with a shorter than average generation time. The corresponding results hold for mother cells generated from daughter cells, and mother cells generated from mother cells. For a sizer model, all daughter cells or mother cells are born at the same average daughter or mother size, irrespective of that cell’s lineage. In this case the positive term vanishes, leaving only the negative correlation arising from the correction of noise-induced fluctuations in cell size as shown in Figure S3.

Figure S3 shows good agreement between our predictions and the correlation coefficients measured in our simulations, both between parent cells and their progeny, and between the two cells generated from a single division event. We observe positive correlations across a broad range of α, x and *σ_t_* values. Using simulations we also investigated the effect of nonvanishing growth rate noise, finding that large *σ_λ_* suppressed these positive correlations, but had little effect for biologically relevant regimes with *σ_λ_/λ*_0_ ≈ 0.15 [7], as shown in Figure S3 (E-F). We also tested the effect of growth morphology, simulating population growth for cells dividing with a budding morphology. Doing so led to additional complexity which is not captured by our theoretical predictions for non-budding cells, and which became more pronounced for increasing *α, σ_t_* and *x* (see Figure S4). These deviations are expected, since the effects of budding on cell division and cell cycle timing are only expected to arise when cells both regulate their size and display variability in cell size (due here to the introduction of division time noise). However, even in the case of budding cells we still find positive generation time correlations across a broad region of parameter space. The authors in Ref. [3] quote a characteristic value of *R*^2^ = 0.25 for the generation time correlation between the mother and daughter cells generated from a cell division event. This aligns well with our our model’s predictions, as shown in Figure S3. These findings motivate a hypothesis that epigenetically inherited division times in budding yeast may arise as a simple consequence of cell size control, without directly affecting the population growth rate as was thought. We also emphasize that although correlated generation times are often a consequence of cell size control, the observation of these correlations in simulated populations of cells is not in itself *sufficient* to generate cell size control.

### 2.5 Growth rate penalty and correlated growth rates

Our findings in Section 2.3 indicate that subject to our model’s assumptions, a single-celled organism is expected to experience a selective pressure to minimize noise in the single cell growth rate. However, for a fixed *σ_λ_* our model predicts that an organism with strong size control might ameliorate its growth rate deficit by dividing asymmetrically. This surprising finding appears inconsistent with the observation of symmetrically dividing cells displaying both size control and noise in their volume growth rates [7], and prompted us to revisit the assumptions underpinning our modeling approach.

To first ensure that our results were robust to minor differences in model structure, we simulated cells following a more detailed inhibitor dilution model. In this model, a stable molecular inhibitor of cell cycle progression must be diluted through growth in order for cells to pass through an essential cell cycle checkpoint (known as Start in the case of budding yeast), and is then newly synthesized prior to cell division. Our prior work investigated the adder correlations that arise within an inhibitor dilution model [20]. Here we use a tunable parameter *a* to describe the degradation of some fraction of a cell’s stock of inhibitor once the cell has passed through Start. By tuning *a* between 0 and 1, this can vary the strategy of size control between a sizer at a = 1 in which all a cell’s inhibitor is completely degraded and newly synthesized with each cell cycle, and an adder at a = 0 in which no degradation takes place but inhibitor synthesis still occurs (see Section S5 for details). The sizer case of this model with *a* = 1 shows good agreement with Equation 5 (see Figure S1 (C)), while the qualitative behavior of decreasing growth rate for weaker size control strategies reproduces that obtained using Equation 4.

We also tested whether this inhibitor dilution model was able to generate robust positive correlations between the generation times of closely related cells, finding that despite quantitative differences arising from differences in model structure, the qualitative findings of Section 2.4 remain intact in this case, as shown in Figure S4 (E,F).

As a further confirmation of our approach for non-IGT size control strategies, we numerically solved Equation 3 for Λ_*P*_ based on the distribution *f*_0_ (*τ*) generated by our simulations (Section S6), and compared our results to the direct fitting of Λ_*P*_ based on the population growth over time. These results are plotted in Figure S5 and show strong agreement between these two approaches.

#### 2.5.1 Non-exponential growth

Experimental evidence demonstrates that excessively large budding yeast cells (≥ 200*fL*, relative to a population average size of ≈ 50*fL*) are known to deviate from exponential growth [21, 22]. This observation has also been predicted on theoretical grounds, due to a low DNA concentration becoming rate-limiting for transcription in excessively large cells [23]. Similarly, excessively small cells are also expected to suffer a fitness cost in their growth rate (for example, due to a limiting abundance of resources for essential cell functions). Motivated by these results, we explored the impact on the population growth rate of a growth rate penalty for cells whose volume deviates from some “optimal” value (see Section S7 for details). Within a biologically relevant range for x, Figure S6 shows a significant decrease in Λ_*P*_ with increasing division asymmetry *x* for cells with weak size control strategies for the parameters tested here. This result is intuitive since broad size distributions will be more penalized by a given growth rate penalty. This finding therefore highlights the need for further experiments that investigate the connection between average cell size and population growth rate, in order to place constraints on the magnitude of such a growth penalty.

#### 2.5.2 Correlated growth rates

To investigate the effect of correlated single-cell growth rates *λ*, we used a model in which the Pearson correlation coefficient (PCC) in *λ* between a parent cell and its progeny could be varied systematically [2] (see Section S8 for details). We tuned the PCC between 0 and 1, finding two qualitatively different regimes for the behavior of Λ_*P*_. Figure S7 demonstrates that for a PCC below 0.5, the effect of growth-rate correlations is minimal, with similar qualitative behavior to that presented in Figure 2. In contrast, large growth rate correlations ≥ 0.5 alter the effect of growth rate noise on Λ_*P*_, leading to an *increase* in Λ_*P*_ with increasing *σ_λ_*, consistent with previous results [10]. Figure S7 further shows that within this regime of strong correlations, increasing division asymmetry *negatively* affects the population growth rate. Experimental observations in *E. coli* show weak correlations in the single cell growth rate with a PCC between mother and daughter growth rates of less than 0.1, indicating that the results of Figure 2 are expected to hold in this case [2, 24].

## 3 Discussion

We study the population growth rate Λ_*P*_ of asymmetrically dividing cells, obtaining analytic expressions for Λ_*P*_ which were confirmed by comparison with simulations. We find that the population growth rate for cells that regulate their size is primarily determined by the single cell growth rate λ, and demonstrate that stochasticity in λ decreases Λ_*P*_ for a model in which noise is coarse-grained over the full cell cycle. This finding is consistent with recent work on this subject in the context of symmetric cell division [2], but conflicts with the interpretation of other studies which predicted that increased noise in generation times will enhance Λ_*P*_, based on models that do not incorporate cell size control [9, 3]. One study presented analytical arguments to support the conclusion that the population doubling time is consistently lower than the average single cell doubling time: (〈*t_d_*) — *T_D_*)/*T_D_* ≥ 0 [9]. Indeed, this inequality is still expected to hold in the case of an asymmetrically dividing population.

This may readily be seen by simple application of Jensen’s inequality to the average of the convex function 〈*e*^-Λ*_p_t*^〉 within the Euler-Lotka equation. Within the class of models we study, the observation that the population doubling time is smaller than the average single cell doubling time does not imply that stochasticity enhances the population growth rate.

Our model further predicts that cells with strong cell size regulation can offset the growth rate deficit that noise in λ generates by dividing asymmetrically. To our knowledge, this is the first model in which exponentially growing cells display a population growth rate that depends on the strategy of size control. Ideally, the predictions we have made here would be tested experimentally by directly varying x for cells with strong size regulation and testing the population growth rate.

To reconcile our results with the abundance of symmetrically dividing organisms throughout nature, we point out that there are many possible scenarios regarding the strength of selection for a higher population growth rate. In one scenario, rapid population growth is the most strongly selected parameter in evolution over many microbial lifecycles, in which case we must conclude that some biologically relevant feature is not incorporated in our model since asymmetric division is clearly not as prolific as would be expected. In another scenario, some organisms, such as yeasts, are subject to occasional strong selection for rapid growth which may lead to asymmetric division based on the predictions we have made here, while for most organisms selection for rapid growth is less important compared to other selections such as survival in harsh environments. In this second scenario, there may be tradeoff costs to asymmetric division that are not evident in exponential growth, causing symmetric division to be favored. Indeed, recent work has demonstrated the existence of a universal tradeoff between the population growth rate and the lag time in bacteria, emphasizing that the population growth rate is not the sole parameter under selection in a given growth medium [25].

Within the first scenario, it may be the case that asymmetric division is difficult to achieve without compromising other aspects of bacterial growth (e.g. by increasing noise in the single cell growth rate), thereby preventing symmetrically dividing organisms from taking advantage of the associated growth rate gains. Another possibility is that the underlying assumptions in our model are flawed. One such assumption was that single-cell growth is truly exponential and independent of cell size. A modified version of our model included a growth rate penalty for excessively large or small cells. When we increased the size of this penalty it eliminated the growth-rate gains associated with asymmetric division, with weaker size control strategies experiencing a more severe growth rate penalty. This result highlights the importance of measuring the variation in growth rate as cell volume deviates from the population average, to further our understanding of the potential advantages and disadvantages of different size control strategies in constraining the spread in cell size. Interest in this area has risen in recent years due to the widespread observations of adder size control strategies in a range of organisms [7, 8, 15], and recent experimental work has made significant steps towards this goal in budding yeast [22], but more work is needed.

One related hypothesis is that asymmetrically dividing cells in nature exist at a local maximum in Λ_*P*_ resulting from a balance between the aforementioned growth rate gains of asymmetric division and a growth rate penalty for unusually sized cells (Figure S6). If this hypothesis is true, our model makes the intuitive prediction that weakening size control substantially in asymmetrically dividing cells without adversely affecting other physiological parameters will lead to a decrease in the population growth rate.

Other simplifying assumptions in our model may also warrant consideration. We assumed that both both cell types in an asymmetrically dividing population will follow the same size control strategy, however, in budding yeast this is not the case, with mother cells displaying weaker size control than daughter cells [8]. A further assumption is that the division rate does not become limited by essential cell cycle processes. This assumption is expected to break down once larger mother cells divide rapidly enough to be limited by the replication and segregation of chromosomes.

We found that positive generation time correlations can be generated by cell size control in asymmetrically dividing cells, contrasting with the negative generation time correlations predicted by the same model for symmetrically dividing cells. This finding motivates a hypothesis for the origin of experimentally observed epigenetic inheritance of division times in closely related budding yeast cells [3].

Our prior work found a significant effect of cell growth geometry on the success of a cell’s size control strategy, predicting that within a budding growth morphology, size control is necessarily ineffective for symmetrically dividing cells [20]. Our collective results here further highlight the importance of studying the effects of different cell growth morphologies, demonstrating that even in the context of exponentially growing cells, asymmetric division can lead to unexpected and novel results. Given the range of diverse growth morphologies that are still being discovered, this demonstrates the need for further investigation of the physiological effects that can arise from novel growth morphologies [1].

## 4 Acknowledgements and Funding Sources

F. B. was supported by the William Georgetti Trust, a Harvard Graduate Merit Award and a Harvard Quantitative Biology Initiative Student Award while conducting this research. A. W. M. thanks NIH grant RO1-GM43987 and the NSF-Simons Center for Mathematical and Statistical Analysis of Biology at Harvard (NSF #1764269, Simons #594596) for support. A. A. acknowledges the support of the NSF CAREER award number 1752024 and the support of the Volkswagen Foundation.

The authors would also like to thank Ethan Levien and Jie Lin for helpful discussions and observations.

## 5 Methods

All simulations of population growth were done using custom-designed code. Our simulations used discretized timesteps to track population growth, and each condition was repeated at least 100 times to generate accurate statistical averages. Populations were seeded with an asynchronous population of 100 cells in equal numbers of cell types D and M, then allowed to propagate for 3.5 population doubling times. Cells were then randomly selected from this population and used to re-seed a new simulation that ran for 6 population doubling times. This was done to maximize the attainment of a steady state generation time distribution. The growth of this reseeded population was then used to infer the population growth rate. All relevant code and Mathematica scripts are available online at https://github.com/felixbarber/division_asymmetry_growth_rate_simulations.

To infer Λ_*p*_ from our simulations, one may measure the growth rate directly based on cell number, or based on total population volume. As noted in [2], these values are identical for cells that display size regulation and therefore have a constant average volume 〈*V*〉(*t*) = Σ_cells_ *V_i_*(*t*))/*N*(*t*) = 〈*V*〉 at steady state. Since the population volume grows continuously and is readily measured in our simulations, the volume growth rate may be more accurately calculated than the number growth rate [2]. We therefore inferred Λ_*p*_ based on measurements of the population volume growth rate throughout this text.

Section S7 explored the behavior of a cell size-dependent average volume growth rate. To ensure a non-negative growth rate for exceptionally large or small cells that were simulated according to this growth policy, whenever a cell was generated with a negative growth rate we removed that cell from consideration.

