## Supplementary Information for "Modeling the impact of single-cell stochasticity and size control on the population growth rate in asymmetrically dividing cells"

November 30, 2020

1: Department of Molecular and Cellular Biology, Harvard University, Cambridge, MA 02138, USA
2: FAS Center for Systems Biology, Harvard University, Cambridge, MA 02138, USA
3: School of Engineering and Applied Sciences, Harvard University, Cambridge, MA 02138, USA

#### Contents

**1 Growth of coupled cell populations** **1**

**2 Two population Euler-Lotka equation** **2**

**3 Approximate solutions to the Euler-Lotka equation** **4**

**4 Non-IGT Perturbation Theory** **6**
4.1 Perturbation theory for $\alpha \approx 1$ . . . . . . . . . . 7

**5 Tunable Inhibitor Dilution Model** **10**

**6 Testing the Euler-Lotka equation for non-IGT growth policies** **11**

**7 Growth rate penalty model** **11**

**8 Growth rate correlations** **12**

**9 Positive generation time correlations** **12**
9.1 Calculations of Generation Time Correlations . . . . . . . . . . 13

**10 Supplementary Figures** **16**

### 1 Growth of coupled cell populations

Here we derive the exponential growth of a population of asymmetrically dividing cells, starting with Equation system 2 in the main text. We assume that we have reached a steady state composition of the population, such that the fractional difference in population size $m(t) = (N_D(t) - N_M(t))/(N_D(t) + N_M(t))$ is constant. We therefore obtain

$$\begin{aligned}\frac{\mathrm{d}N_D}{\mathrm{d}t} &= \Lambda_M \frac{1-m}{1+m} N_D(t),\\ \frac{\mathrm{d}N_M}{\mathrm{d}t} &= \Lambda_D \frac{1+m}{1-m} N_M(t).\end{aligned} \tag{1}$$

These equations imply exponential growth with growth rates $\Lambda_M \frac{1-m}{1+m}$ and $\Lambda_D \frac{1+m}{1-m}$. In order for this to yield a constant fraction $m$ we require that these growth rates be equal, implying $\Lambda_D/\Lambda_M = \frac{(1-m)^2}{(1+m)^2}$. If we now consider $N(t) = N_D(t) + N_M(t)$, then the above implies

$$\frac{\mathrm{d}N}{\mathrm{d}t} = \Lambda_P N(t), \tag{2}$$

where $\Lambda_P = \sqrt{\Lambda_D \Lambda_M}$ as stated in the main text.

### 2 Two population Euler-Lotka equation

In this section we determine the corresponding Euler-Lotka equation for an asymmetrically dividing population, showing that it has the same form as Equation 1 in the main text. We also obtain a second constraint that allows one to calculate the population asymmetry $m$ defined in Section 1. In doing so we follow a similar approach to [1].

Given a distribution of generation times $f_0^D(\gamma)$ for cell type $D$, we define the survival function $F_-^D(\gamma) = \int_\gamma^\infty f_0^D(y)\mathrm{d}y$ to be the probability of a cell of that type surviving until age $\gamma$. We then define the division rate $\mu^D(\gamma)$ for cells of type $D$ and age $\gamma$ as

$$\begin{aligned}\mu^D(\gamma) &= \frac{F_-^D(\gamma) - F_-^D(\gamma + \mathrm{d}\gamma)}{F_-^D(\gamma)\mathrm{d}\gamma}\\ &= \frac{f_0^D(\gamma)}{F_-^D(\gamma)}\end{aligned} \tag{3}$$

Note that $f_0(\gamma)$ is defined as the distribution of generation times for a cell of a given type that has just been born in a given small window of time. This definition also corresponds to the distribution of generation times measured over the full lineage tree for that cell type [2], which is more convenient in practice. This can be seen by splitting the time during which a population is growing into infinitessimal time slices and considering only those cells have been born in each slice. Doing so we see that provided the population has reached steady state, the generation times of all cells sampled in this way will be drawn from $f_0(\gamma)$. By integrating over all slices, we can sample each cell in the complete history of the population, and observe that the distribution of generation times measured over the full population must also be $f_0(\gamma)$. The distribution of generation times measured over the full population is therefore the same as that for a cell that has just been born. Within our model the birth generation time distribution $f_0(\gamma)$ is the meaningful distribution with which to calculate the division rate $\mu$ for a randomly sampled cell of age $\gamma$. Since this is equivalent to the full tree distribution, this is what we use to calculate $\mu(\gamma)$.
We also define $\Lambda_D = \int_0^\infty \phi^D(\gamma)\mu^D(\gamma)\mathrm{d}\gamma$ as the average rate of division per daughter cell, where $\phi^D(\gamma)$ is the age distribution of cells of type $D$, and $\mu^D(\gamma)$ is the rate of division for cells of type $D$ and age $\gamma$. Similar definitions hold for cell type $M$.
Given the above definitions, the following relation holds:

$$\begin{aligned}N_D(t)\phi^D(\gamma)(1 - \mu^D(\gamma)\mathrm{d}t) &= \phi^D(\gamma + \mathrm{d}t)N_D(t + \mathrm{d}t)\\ &= (\phi^D(\gamma) + \mathrm{d}t\frac{\mathrm{d}\phi^D}{\mathrm{d}t})(N_D(t) + \Lambda_M N_M(t)\mathrm{d}t).\end{aligned} \tag{4}$$

We therefore obtain

$$\begin{aligned}\frac{\mathrm{d}\phi^D}{\mathrm{d}t} &= -\phi^D(\gamma)\left(\mu^D(\gamma) + \Lambda_M \frac{N_M(t)}{N_D(t)}\right) \\ &= -\phi^D(\gamma)(\mu^D(\gamma) + \Lambda_P) \\ \Rightarrow \phi^D(\gamma) &= \phi_0^D \exp\left(-\Lambda_P\gamma - \int_0^\gamma \mu^D(y)\mathrm{d}y\right) \\ &= \phi_0^D \exp\left(-\Lambda_P\gamma\right) F_-^D(\gamma)\end{aligned} \tag{5}$$

The last line is obtained by noting that $f_0^D(\gamma) = -\mathrm{d}F_-^D(\gamma)/\mathrm{d}\gamma$ , so that Equation S3 implies $F_-^D(\gamma) = \exp\left(-\int_0^\gamma \mu^D(y)\mathrm{d}y\right)$. Normalization of $\phi^D$ requires that

$$1 = \int_0^\infty \phi^D(\gamma)\mathrm{d}\gamma.$$

Integrating by parts yields

$$\begin{aligned}1 &= \phi_0^D \frac{1}{\Lambda_P}\left(1 - \int_0^\infty f_0^D(\gamma)\exp(-\Lambda_P\gamma)\mathrm{d}\gamma\right) \\ &= \frac{1}{\Lambda_P}(\phi_0^D - \Lambda_D) \\ \Rightarrow \phi_0^D &= \Lambda_P(1 + \sqrt{\Lambda_D/\Lambda_M}).\end{aligned} \tag{6}$$

We now calculate the PDF $f_1(\tau)$ for branch cells (all previous cells in the distribution at a given point in time) that divide at age $\tau$.

$$\begin{aligned}f_1^D(\tau) &= \frac{N_D(t)\phi^D(\tau)\mu^D(\tau)\mathrm{d}\tau}{N_D(t)\Lambda_D\mathrm{d}\tau} \\ &= (1 + \sqrt{\frac{\Lambda_M}{\Lambda_D}})\mu^D(\tau)\exp\left(-\Lambda_P\tau - \int_0^\tau \mu^D(\gamma)\mathrm{d}\gamma\right) \\ &= (1 + \sqrt{\frac{\Lambda_M}{\Lambda_D}})\exp\left(-\Lambda_P\tau\right) f_0^D(\tau)\end{aligned} \tag{7}$$

The numerator in the first line is the number of cells that divide at age $\tau$ in time $\mathrm{d}t$, while the denominator is the overall number of divisions in this time. Integrating both sides yields the normalization condition

$$1 = (1 + \sqrt{\frac{\Lambda_M}{\Lambda_D}})\int_0^\infty \exp\left(-\Lambda_P\tau\right) f_0^D(\tau)\mathrm{d}\tau. \tag{8}$$

Through the same process as above we also obtain the corresponding condition for $f_1^M(\tau)$

$$1 = (1 + \sqrt{\frac{\Lambda_D}{\Lambda_M}})\int_0^\infty \exp\left(-\Lambda_P\tau\right) f_0^M(\tau)\mathrm{d}\tau. \tag{9}$$

We now recall that for $-1 < m < 1$, $\sqrt{\frac{\Lambda_D}{\Lambda_M}} = (1-m)/(1+m)$. Rearranging Equations S8 and S9 to eliminate $m$ yields Equation 1 from the main text, restated here:

$$1 = 2\int_0^\infty \exp\left(-\Lambda_P\tau\right) f_0^P(\tau)\mathrm{d}\tau, \tag{10}$$

where $f_0^P(\tau) = \frac{1}{2}(f_0^D(\tau) + f_0^M(\tau))$. We see therefore that the Euler-Lotka equation holds for the two cell type case, by taking the distribution $f_0^P$ to be the full tree distribution measured for both cell types. We note that although there may be an imbalance in the number of cells of different types present at a given point in time as indicated by $m$, over the full history of the population this imbalance will be removed, and there will be approximately equal numbers of each cell type, leading to the factor of $1/2$ in Equation S10. We also have an equation determining $m$ as

$$m = \int_0^\infty \exp\left(-\Lambda_P \tau\right) \left(f_0^M(\tau) - f_0^D(\tau)\right) \mathrm{d}\tau. \tag{11}$$

Equation S10 therefore determines the population growth rate $\Lambda_P$, which may then be inserted into Equation S11 to determine the ratio of the two cell-types $m$.

### 3 Approximate solutions to the Euler-Lotka equation

We first consider the special case of a timer for cell division, without noise in generation times or in the single cell growth rate, i.e. $f_0^D(\tau) = \delta(\tau - \tau_D)$ and $f_0^M(\tau) = \delta(\tau - \tau_M)$. In this case, Equation S10 simplifies to the implicit equation for $\Lambda_P$

$$\exp(-\Lambda_P \tau_D) + \exp(-\Lambda_P \tau_M) = 1. \tag{12}$$

This case describes the non-stochastic model of [3]. The result in Equation S12 was originally derived in [4] by different means.

We now consider a sizer model ($\alpha = 1$), neglecting noise in interdivision times but allowing noise in the growth rate $\lambda$. In this limit, the cell volume at division is approximately $2\Delta$ regardless of celltype, so that the volume at birth is $(1-x)\Delta$ for daughters and $(1+x)\Delta$ for mothers. Since the volume at birth is independent of the previous cell cycle, this growth policy is well described by the IGT model, allowing us to calculate an approximate analytic expression for the population growth rate. Within this model the generation times are $t_d = \ln(2/(1-x))/\lambda$ for cell type $D$ and $t_d = \ln(2/(1+x))/\lambda$ for cell type $M$. We proceed by treating the two integrals in Equation S10 separately. For cell type $M$, reparameterizing Equation S8 by $\omega = \ln((1+x)/2)/\tau$ implies

$$\begin{aligned}
&\int_0^\infty \exp\left(-\Lambda_P \tau\right) f_0^M(\tau) \mathrm{d}\tau = \\
&\quad \frac{1}{\sqrt{2\pi\sigma_\lambda^2}} \int_0^\infty \exp\left[-\Lambda_P \ln(2/(1+x))/\omega\right] \exp\left[-\frac{(\omega-\lambda_0)^2}{2\sigma_\lambda^2}\right] \mathrm{d}\omega, \\
&\quad = \frac{1}{\sqrt{2\pi\sigma_\lambda^2}} \int_0^\infty \exp\left[-\frac{1}{2\sigma_\lambda^2}\left(2\sigma_\lambda^2 \Lambda_P \ln(2/(1+x))/\omega + (\omega-\lambda_0)^2\right)\right] \mathrm{d}\omega, \\
&\quad = \frac{1}{\sqrt{2\pi\sigma_\lambda^2}} I_M,
\end{aligned} \tag{13}$$

where we define

$$\begin{aligned}
I_M &= \int_0^\infty \exp\left[-\frac{1}{2\sigma_\lambda^2}\left(2\sigma_\lambda^2 \Lambda_P \ln(2/(1+x))/\omega + (\omega-\lambda_0)^2\right)\right] \mathrm{d}\omega \\
&= \int_0^\infty \exp\left[-\frac{1}{2\sigma_\lambda^2} g_M(\omega)\right] \mathrm{d}\omega.
\end{aligned} \tag{14}$$

$I_M$ may be approximated using the saddle point method

$$I_M = \exp\left[-\frac{1}{2\sigma_\lambda^2} g_M(\omega_{c,M})\right] \sqrt{\frac{4\pi\sigma_\lambda^2}{g_M''(\omega_{c,M})}}, \tag{15}$$

where $\omega_{c,M}$ is defined by $g'_M(\omega_{c,M}) = 0$ with $g''_M(\omega_{c,M}) > 0$. This condition yields the expressions

$$\begin{aligned}
\omega_{c,M} &= \lambda_0 - \frac{\Lambda_P \sigma_\lambda^2 \ln((1+x)/2)}{\lambda_0^2}, \\
g_M(\omega_{c,M}) &= \frac{\Lambda_P^2 \sigma_\lambda^4 \left(\ln(\frac{1+x}{2})\right)^2}{\lambda_0^4} + \frac{2\Lambda_P {\sigma_\lambda}^2 \ln\left(\frac{2}{1+x}\right)}{\lambda_0 - \frac{\Lambda_P {\sigma_\lambda}^2 \ln(\frac{1+x}{2})}{\lambda_0^2}}, \\
g''_M(\omega_{c,M}) &= \frac{4\Lambda_P {\sigma_\lambda}^2 \ln\left(\frac{2}{1+x}\right)}{\left(\lambda_0 - \frac{\Lambda_P {\sigma_\lambda}^2 \ln((1+x)/2)}{\lambda_0^2}\right)^3} + 2.
\end{aligned} \tag{16}$$

Performing similar analysis to the above for the second integral in Equation S10 yields the corresponding relations for cell type $D$

$$\begin{aligned}
\omega_{c,D} &= \lambda_0 - \frac{\Lambda_P \sigma_\lambda^2 \ln(\frac{1-x}{2})}{\lambda_0^2}, \\
g_D(\omega_{c,D}) &= \frac{\Lambda_P^2 \sigma_\lambda^4 \left(\ln(\frac{1-x}{2})\right)^2}{\lambda_0^4} + \frac{2\Lambda_P {\sigma_\lambda}^2 \ln\left(\frac{2}{1-x}\right)}{\lambda_0 - \frac{\Lambda_P {\sigma_\lambda}^2 \ln(\frac{1-x}{2})}{\lambda_0^2}}, \\
g''_D(\omega_{c,D}) &= \frac{4\Lambda_P {\sigma_\lambda}^2 \ln\left(\frac{2}{1-x}\right)}{\left(\lambda_0 - \frac{\Lambda_P {\sigma_\lambda}^2 \ln(\frac{1-x}{2})}{\lambda_0^2}\right)^3} + 2.
\end{aligned} \tag{17}$$

We substitute the saddle point approximation for $I_M$ and $I_D$ into Equation S10. We proceed to Taylor expand the right hand sides to $2^{nd}$ order in $\Lambda_P$ and to $4^{th}$ order in $\sigma_\lambda$. Solving for $\Lambda_P$ while keeping only the lowest order terms in $\sigma_\lambda$ gives

$$\Lambda_P(\sigma_\lambda, x) = \lambda_0 \left(1 - \left(1 + \frac{1}{2} \frac{(1+x)(\ln(\frac{1+x}{2}))^2 + (1-x)(\ln(\frac{1-x}{2}))^2}{(1+x)\ln(\frac{1+x}{2}) + (1-x)\ln(\frac{1-x}{2})}\right)\left(\frac{\sigma_\lambda}{\lambda_0}\right)^2\right) + O(\sigma_\lambda^4). \tag{18}$$

Setting $x = 0$ in Equation S18 recovers the solution from [1] for symmetric growth of

$$\lambda_p(\sigma_\lambda) = \lambda_o \left(1 - \left(1 - \frac{\ln 2}{2}\right)\left(\frac{\sigma_\lambda}{\lambda_o}\right)^2\right). \tag{19}$$

Inserting the result for $\Lambda_P$ from Equation S18 to second order in $\sigma_\lambda$ into Equation S11 for the population asymmetry $m$, we obtain the satisfyingly simple equality $m = x$. This result is tested in Figure S2 (A), showing excellent agreement in the case of a sizer for varying levels of $\sigma_\lambda$. The increase in $m$ with $x$ is intuitive, since increasing the division asymmetry between mother and daughter cells leads to a larger discrepancy in their sizes and consequently their generation times, consequently leading to a greater bias for the longer lived daughter cells in a snapshot of the population. Our simulations further demonstrate that $m$ is independent of $\sigma_\lambda$ for other strategies of size control (see Figure S2 (A)). Figure S2 (B) demonstrates the behavior of $m$ for different size control strategies, showing that weaker size control strategies decrease the asymmetry $m$. This decrease in $m$ with decreasing $\alpha$ is also intuitive, since weaker size control strategies decrease the discrepancy in generation times between the larger mother and smaller daughter cells, decreasing the bias towards the longer lived daughter cells present in a snapshot of the population.

### 4 Non-IGT Perturbation Theory

In this section we derive the perturbative result for small $1-\alpha$ shown in Figure 2C in the main text. Because the volume is always a positive value, we will use the log of volume as a variable: $X_b = \ln(V_b/\Delta)$. Then we shall define a tree distribution of $X_b$ for daughters ($\psi_D[X_b]$) and mothers ($\psi_M[X_b]$). Using the transport equation approach [2], we can write down a self-consistent equation for each distribution.

$$\psi_D[X_b] = \int\int \psi[X_b']p[X_b,\tau,D|X_b']e^{-\Lambda_p\tau}dX_b'd\tau \tag{20}$$

$$\psi_M[X_b] = \int\int \psi[X_b']p[X_b,M,\tau|X_b']e^{-\Lambda_p\tau}dX_b'd\tau, \tag{21}$$

where the transition probabilities are determined by our growth model:

$$\begin{aligned} p[X_b,\tau,D|X_b']dX_b'd\tau = \int \delta(X_b - \ln(1-x) - \ln(\alpha+(1-\alpha)e^{X_b'})\Delta(1-x)(\alpha+(1-\alpha)e^{X_b'}) \\ \times\left(\tau - \frac{1}{\lambda}\ln\left(\frac{2\alpha}{e^{X_b'}} + 2(1-\alpha)\right)\right) f(\lambda)dX_b'd\tau d\lambda, \end{aligned} \tag{22}$$

$$\begin{aligned} p[X_b,\tau,M|X_b']dX_b'd\tau = \int \delta(X_b - \ln(1+x) - \ln(\alpha+(1-\alpha)e^{X_b'})\Delta(1-x)(\alpha+(1-\alpha)e^{X_b'}) \\ \times\left(\tau - \frac{1}{\lambda}\ln\left(\frac{2\alpha}{e^{X_b'}} + 2(1-\alpha)\right)\right) f(\lambda)dV_b'd\tau d\lambda. \end{aligned} \tag{23}$$

We expect the average $X_b$ for the mother cells to be about $\ln(1+x)$ and for the daughter cells $\ln(1-x)$. Defining $\delta X_b = X_b - \ln(1\pm x)$, and since Equations 22 and 23 differ only in the term $\ln(1\pm x)$ we find the average of $F[\delta X_b]$, where $F$ is some arbitrary function, to be the same for the mother and daughter:

$$\begin{aligned} \int F\left[X_b - \ln(1-x)\right]\psi_D[X_b]dX_b \equiv \langle \delta F[\delta X_b]\rangle_D, \\ \int F\left[X_b - \ln(1+x)\right]\psi_M[X_b]dX_b \equiv \langle \delta F[\delta X_b]\rangle_M, \end{aligned} \tag{24}$$

$$\begin{aligned} \langle F[\delta X_b]\rangle_D &= \langle F[\delta X_b]\rangle_M \\ &= \int F\left[\ln(\alpha+(1-\alpha)e^{X_b'})\right]\psi[X_b']f(\lambda)\exp\left[-\frac{\Lambda_p}{\lambda}\ln\left(\frac{2\alpha}{e^{X_b'}} + 2(1-\alpha)\right)\right]dX_b'd\lambda. \end{aligned} \tag{25}$$

By using $F[\delta X_b] = 1$, we get

$$\begin{aligned} \int \psi_D[X_b]dX_b = \frac{1}{2} \\ &= \int \psi[X_b']f(\lambda)\exp\left[-\frac{\Lambda_p}{\lambda}\ln\left(\frac{2\alpha}{e^{X_b'}} + 2(1-\alpha)\right)\right]dX_b'd\lambda \\ &= \left\langle \int f(\lambda)\exp\left[-\frac{\Lambda_p}{\lambda}\ln\left(\frac{2\alpha}{(1-x)e^{\delta X_b'}} + 2(1-\alpha)\right)\right]d\lambda\right\rangle_D \qquad (26) \\ &+ \left\langle \int f(\lambda)\exp\left[-\frac{\Lambda_p}{\lambda}\ln\left(\frac{2\alpha}{(1+x)e^{\delta X_b'}} + 2(1-\alpha)\right)\right]d\lambda\right\rangle_M. \qquad (27) \end{aligned}$$

The first term can be approximated by series expansion and we define $A = (\Lambda_P - \lambda_0)/\lambda_0$.

$$\left\langle \int f(\lambda) \exp\left[ -\frac{\Lambda_p}{\lambda} \ln\left( \frac{2\alpha}{(1-x)e^{\delta X_b'}} + 2(1-\alpha) \right) \right] d\lambda \right\rangle_D \tag{28}$$

$$\approx \left\langle \int f(\lambda) \exp\left[ -(1+A) \left( 1 - \frac{\lambda - \lambda_0}{\lambda_0} + \frac{(\lambda-\lambda_0)^2}{\lambda_0^2} \right) \right. \right. \tag{29}$$

$$\left. \left. \times \left( \ln\left( \frac{2-2(1-\alpha)x}{1-x} \right) + \ln\left( \frac{1-x}{1-(1-\alpha)x} \frac{\alpha + (1-\alpha)(1-x)e^{\delta X_b'}}{e^{\delta X_b'}(1-x)} \right) \right) \right] d\lambda \right\rangle_D . \tag{30}$$

We simplify the equation by assuming that $\ln\left( \frac{1-x}{1-(1-\alpha)x} \frac{\alpha+(1-\alpha)(1-x)e^{\delta X_b'}}{e^{\delta X_b'}(1-x)} \right)$ is small as long as $\alpha$ is close to 1 and the asymmetry is small.

$$\begin{aligned}
&\left\langle \int f(\lambda) \exp\left[ -\frac{\Lambda_p}{\lambda} \ln\left( \frac{2\alpha}{(1-x)e^{\delta X_b'}} + 2(1-\alpha) \right) \right] d\lambda \right\rangle_D \\
&= \left\langle \int f(\lambda) \frac{1-x}{2-2(1-\alpha)x} \exp\left[ \left( -A + \frac{\lambda-\lambda_0}{\lambda_0} - \frac{(\lambda-\lambda_0)^2}{\lambda_0^2} \right) \ln\left( \frac{2-2(1-\alpha)x}{1-x} \right) \right] \right. \\
&\qquad\qquad \left. \times \frac{1-(1-\alpha)x}{1-x} \frac{e^{\delta X_b'}(1-x)}{\alpha + (1-\alpha)(1-x)e^{\delta X_b'}} d\lambda \right\rangle_D \\
&= \frac{1}{2} \left( 1 - \ln\left( \frac{2-2(1-\alpha)x}{1-x} \right) A - \ln\left( \frac{2-2(1-\alpha)x}{1-x} \right) \left( 1 - \frac{1}{2} \ln\left( \frac{2-2(1-\alpha)x}{1-x} \right) \right) \frac{\sigma_\lambda^2}{\lambda_0^2} \right) \left\langle \frac{e^{\delta X_b'}(1-x)}{\alpha + (1-\alpha)(1-x)e^{\delta X_b'}} \right\rangle_D .
\end{aligned} \tag{31}$$

Similarly, for the mother cell,

$$\left\langle \int f(\lambda) \exp\left[ -\frac{\Lambda_p}{\lambda} \ln\left( \frac{2\alpha}{(1+x)e^{\delta X_b'}} + 2(1-\alpha) \right) \right] d\lambda \right\rangle_M \tag{32}$$

$$= \frac{1}{2} \left( 1 - \ln\left( \frac{2+2(1-\alpha)x}{1+x} \right) A - \ln\left( \frac{2+2(1-\alpha)x}{1+x} \right) \left( 1 - \frac{1}{2} \ln\left( \frac{2+2(1-\alpha)x}{1+x} \right) \right) \frac{\sigma_\lambda^2}{\lambda_0^2} \right) \left\langle \frac{e^{\delta X_b'}(1+x)}{\alpha + (1-\alpha)(1+x)e^{\delta X_b'}} \right\rangle_M . \tag{33}$$

Combining the expressions we get,

$$\begin{aligned}
\frac{1}{2} = &\frac{1}{2} \left( 1 - \ln\left( \frac{2-2(1-\alpha)x}{1-x} \right) A - \ln\left( \frac{2-2(1-\alpha)x}{1-x} \right) \left( 1 - \frac{1}{2} \ln\left( \frac{2-2(1-\alpha)x}{1-x} \right) \right) \frac{\sigma_\lambda^2}{\lambda_0^2} \right) \left\langle \frac{e^{\delta X_b'}(1-x)}{\alpha + (1-\alpha)(1-x)e^{\delta X_b'}} \right\rangle_D \\
&+ \frac{1}{2} \left( 1 - \ln\left( \frac{2+2(1-\alpha)x}{1+x} \right) A - \ln\left( \frac{2+2(1-\alpha)x}{1+x} \right) \left( 1 - \frac{1}{2} \ln\left( \frac{2+2(1-\alpha)x}{1+x} \right) \right) \frac{\sigma_\lambda^2}{\lambda_0^2} \right) \left\langle \frac{e^{\delta X_b'}(1+x)}{\alpha + (1-\alpha)(1+x)e^{\delta X_b'}} \right\rangle_M .
\end{aligned} \tag{34}$$

#### 4.1 Perturbation theory for $\alpha \approx 1$

Let's define $\delta\alpha \equiv 1 - \alpha$ and expand Equation S34 to $O(\delta\alpha^n)$.

$$\frac{1}{2}=\frac{1}{2}\left[1-\left(\ln\left(\frac{2}{1-x}\right)-\sum_{m=1}^{n}\frac{x^m}{m}\delta\alpha^m\right)A-\left(\ln\left(\frac{2}{1-x}\right)-\sum_{m=1}^{n}\frac{x^m}{m}\delta\alpha^m\right)\right.$$
$$\left.\times\left(1-\frac{1}{2}\left(\ln\left(\frac{2}{1-x}\right)-\sum_{m=1}^{n}\frac{x^m}{m}\delta\alpha^m\right)\right)\frac{\sigma_\lambda^2}{\lambda_0^2}\right]\left\langle\sum_{m=0}^{n}(1-(1-x)e^{\delta X_b'})^m(1-x)e^{\delta X_b'}\delta\alpha^m\right\rangle_D$$
$$+\frac{1}{2}\left[1-\left(\ln\left(\frac{2}{1+x}\right)-\sum_{m=1}^{n}\frac{(-x)^m}{m}\delta\alpha^m\right)A-\left(\ln\left(\frac{2}{1+x}\right)-\sum_{m=1}^{n}\frac{(-x)^m}{m}\delta\alpha^m\right)\right.$$
$$\left.\times\left(1-\frac{1}{2}\left(\ln\left(\frac{2}{1+x}\right)-\sum_{m=1}^{n}\frac{(-x)^m}{m}\delta\alpha^m\right)\right)\frac{\sigma_\lambda^2}{\lambda_0^2}\right]\left\langle\sum_{m=0}^{n}(1-(1+x)e^{\delta X_b'})^m(1+x)e^{\delta X_b'}\delta\alpha^m\right\rangle_M+O(\delta\alpha^{n+1}). \tag{35}$$

We can do the same for Equation S25.

$$\left\langle F\left[X_b'\right]\right\rangle_D$$
$$=\frac{1}{2}\left[1-\left(\ln\left(\frac{2}{1-x}\right)-\sum_{m=1}^{n}\frac{x^m}{m}\delta\alpha^m\right)A-\left(\ln\left(\frac{2}{1-x}\right)-\sum_{m=1}^{n}\frac{x^m}{m}\delta\alpha^m\right)\right.$$
$$\left.\times\left(1-\frac{1}{2}\left(\ln\left(\frac{2}{1-x}\right)-\sum_{m=1}^{n}\frac{x^m}{m}\delta\alpha^m\right)\right)\frac{\sigma_\lambda^2}{\lambda_0^2}\right]$$
$$\times\left\langle\sum_{l=0}^{\infty}\sum_{m=0}^{n}(1-(1-x)e^{\delta X_b'})^m(1-x)e^{\delta X_b'}\frac{F^{(l)}[0]}{l!}\left(\ln(1+\delta\alpha(-1+(1-x)e^{\delta X_b'}))\right)^l\delta\alpha^m\right\rangle_D$$
$$+\frac{1}{2}\left[1-\left(\ln\left(\frac{2}{1+x}\right)-\sum_{m=1}^{n}\frac{(-x)^m}{m}\delta\alpha^m\right)A-\left(\ln\left(\frac{2}{1+x}\right)-\sum_{m=1}^{n}\frac{(-x)^m}{m}\delta\alpha^m\right)\right.$$
$$\left.\times\left(1-\frac{1}{2}\left(\ln\left(\frac{2}{1+x}\right)-\sum_{m=1}^{n}\frac{(-x)^m}{m}\delta\alpha^m\right)\right)\frac{\sigma_\lambda^2}{\lambda_0^2}\right]$$
$$\times\left\langle\sum_{l=0}^{\infty}\sum_{m=0}^{n}(1-(1+x)e^{\delta X_b'})^m(1+x)e^{\delta X_b'}\frac{F^{(l)}[0]}{l!}\left(\ln(1+\delta\alpha(-1+(1+x)e^{\delta X_b'}))\right)^l\delta\alpha^m\right\rangle_M+O(\delta\alpha^{n+1}). \tag{36}$$

Here we can find the correction terms for $A=\sum_{i=0}^{n}\delta\alpha^i A_i$, where

$$A_0=\left(-1+\frac{1}{2}\frac{(1-x)\ln\left(\frac{2}{1-x}\right)^2+(1+x)\ln\left(\frac{2}{1+x}\right)^2}{(1-x)\ln\left(\frac{2}{1-x}\right)+(1+x)\ln\left(\frac{2}{1+x}\right)}\right)\frac{\sigma_\lambda^2}{\lambda_0^2}. \tag{37}$$

Let's start with the first order correction. Equation S34 can be rewritten as

$$
\begin{aligned}
\frac{1}{2} = &\frac{1}{2}\left[1-\left(\ln\left(\frac{2}{1-x}\right)-x\delta\alpha\right)(A_0+A_1\delta\alpha)-\left(\ln\left(\frac{2}{1-x}\right)-x\delta\alpha\right)\left(1-\frac{1}{2}\left(\ln\left(\frac{2}{1-x}\right)-x\delta\alpha\right)\right)\frac{\sigma_\lambda^2}{\lambda_0^2}\right]\\
&\left[\left\langle(1-x)e^{\delta X_b'}\right\rangle_D+\left\langle(1-x)e^{\delta X_b'}-(1-x)^2e^{2\delta X_b'}\right\rangle_D\delta\alpha\right]\\
&+\frac{1}{2}\left[1-\left(\ln\left(\frac{2}{1+x}\right)+x\delta\alpha\right)(A_0+A_1\delta\alpha)-\left(\ln\left(\frac{2}{1+x}\right)+x\delta\alpha\right)\left(1-\frac{1}{2}\left(\ln\left(\frac{2}{1+x}\right)+x\delta\alpha\right)\right)\frac{\sigma_\lambda^2}{\lambda_0^2}\right]\\
&\left[\left\langle(1+x)e^{\delta X_b'}\right\rangle_M+\left\langle(1+x)e^{\delta X_b'}-(1+x)^2e^{2\delta X_b'}\right\rangle_M\delta\alpha\right]\\
&=\left\langle e^{\delta X_b'}\right\rangle_D\\
&+\delta\alpha\left[\frac{1}{2}\left(xA_0-\ln\left(\frac{2}{1-x}\right)A_1-\left(\frac{1}{2}\ln\left(\frac{2}{1-x}\right)x-x\right)\frac{\sigma_\lambda^2}{\lambda_0^2}\right)\left\langle(1-x)e^{\delta X_b'}\right\rangle_D\right.\\
&+\frac{1}{2}\left(-xA_0-\ln\left(\frac{2}{1+x}\right)A_1-\left(-\frac{1}{2}\ln\left(\frac{2}{1+x}\right)x+x\right)\frac{\sigma_\lambda^2}{\lambda_0^2}\right)\left\langle(1+x)e^{\delta X_b'}\right\rangle_M\\
&+\frac{1}{2}\left(1-\ln\left(\frac{2}{1-x}\right)A_0-\ln\left(\frac{2}{1-x}\right)\left(1-\frac{1}{2}\ln\left(\frac{2}{1-x}\right)\right)\frac{\sigma_\lambda^2}{\lambda_0^2}\right)\left\langle(1-x)e^{\delta X_b'}-(1-x)^2e^{2\delta X_b'}\right\rangle_D\\
&\left.+\frac{1}{2}\left(1-\ln\left(\frac{2}{1+x}\right)A_0-\ln\left(\frac{2}{1+x}\right)\left(1-\frac{1}{2}\ln\left(\frac{2}{1+x}\right)\right)\frac{\sigma_\lambda^2}{\lambda_0^2}\right)\left\langle(1+x)e^{\delta X_b'}-(1+x)^2e^{2\delta X_b'}\right\rangle_M\right].
\end{aligned}
\tag{38}
$$

This time we need to calculate $\left\langle e^{\delta X_b'}\right\rangle_D$ to $O(\delta\alpha)$ and $\left\langle e^{2\delta X_b'}\right\rangle_D$ to $O(\delta\alpha^0)$. Using Equation S25, we find

$$\left\langle e^{2\delta X_b'}\right\rangle_D = \frac{1}{2}+O(\delta\alpha), \tag{39}$$

$$
\begin{aligned}
\left\langle e^{\delta X_b'}\right\rangle_D = &\frac{1}{2}-\frac{\delta\alpha}{2}+\delta\alpha\left[\frac{1}{2}\left(1-\ln\left(\frac{2}{1-x}\right)A_0-\ln\left(\frac{2}{1-x}\right)\left(1-\frac{1}{2}\ln\left(\frac{2}{1-x}\right)\right)\frac{\sigma_\lambda^2}{\lambda_0^2}\right)\left\langle e^{2\delta X_b'}(1-x)^2\right\rangle_D\right.\\
&\left.+\frac{1}{2}\left(1-\ln\left(\frac{2}{1+x}\right)A_0-\ln\left(\frac{2}{1+x}\right)\left(1-\frac{1}{2}\ln\left(\frac{2}{1+x}\right)\right)\frac{\sigma_\lambda^2}{\lambda_0^2}\right)\left\langle e^{2\delta X_b'}(1+x)^2\right\rangle_M\right]+O(\delta\alpha^2).
\end{aligned}
\tag{40}
$$

Thus, collecting the first order terms in Equation S38, we can calculate $A_1$.

$$\begin{aligned}
&\left(\frac{1-x}{4}\ln\left(\frac{2}{1-x}\right)+\frac{1+x}{4}\ln\left(\frac{2}{1+x}\right)\right)A_1\\
&=-\frac{1}{2}+\left[\frac{1}{2}\left(1-\ln\left(\frac{2}{1-x}\right)A_0-\ln\left(\frac{2}{1-x}\right)\left(1-\frac{1}{2}\ln\left(\frac{2}{1-x}\right)\right)\frac{\sigma_\lambda^2}{\lambda_0^2}\right)\left\langle e^{2\delta X_b'}(1-x)^2\right\rangle_D\right.\\
&\left.+\frac{1}{2}\left(1-\ln\left(\frac{2}{1+x}\right)A_0-\ln\left(\frac{2}{1+x}\right)\left(1-\frac{1}{2}\ln\left(\frac{2}{1+x}\right)\right)\frac{\sigma_\lambda^2}{\lambda_0^2}\right)\left\langle e^{2\delta X_b'}(1+x)^2\right\rangle_M\right]\\
&+\left[\frac{1}{2}\left(xA_0-\left(\frac{1}{2}\ln\left(\frac{2}{1-x}\right)x-x\right)\frac{\sigma_\lambda^2}{\lambda_0^2}\right)\left\langle(1-x)e^{\delta X_b'}\right\rangle_D\right.\\
&+\frac{1}{2}\left(-xA_0-\left(-\frac{1}{2}\ln\left(\frac{2}{1+x}\right)x+x\right)\frac{\sigma_\lambda^2}{\lambda_0^2}\right)\left\langle(1+x)e^{\delta X_b'}\right\rangle_M\\
&+\frac{1}{2}\left(1-\ln\left(\frac{2}{1-x}\right)A_0-\ln\left(\frac{2}{1-x}\right)\left(1-\frac{1}{2}\ln\left(\frac{2}{1-x}\right)\right)\frac{\sigma_\lambda^2}{\lambda_0^2}\right)\left\langle(1-x)e^{\delta X_b'}-(1-x)^2e^{2\delta X_b'}\right\rangle_D\\
&\left.+\frac{1}{2}\left(1-\ln\left(\frac{2}{1+x}\right)A_0-\ln\left(\frac{2}{1+x}\right)\left(1-\frac{1}{2}\ln\left(\frac{2}{1+x}\right)\right)\frac{\sigma_\lambda^2}{\lambda_0^2}\right)\left\langle(1+x)e^{\delta X_b'}-(1+x)^2e^{2\delta X_b'}\right\rangle_M\right]\\
&=\frac{1-x}{4}\left(xA_0-\left(\frac{1}{2}\ln\left(\frac{2}{1-x}\right)x-x\right)\frac{\sigma_\lambda^2}{\lambda_0^2}\right)+\frac{1+x}{4}\left(-xA_0-\left(-\frac{1}{2}\ln\left(\frac{2}{1+x}\right)x+x\right)\frac{\sigma_\lambda^2}{\lambda_0^2}\right)\\
&=-\frac{x^2}{2}A_0-\left(\frac{x^2}{2}+\frac{(1-x)x}{8}\ln\left(\frac{2}{1-x}\right)-\frac{(1+x)x}{8}\ln\left(\frac{2}{1+x}\right)\right)\frac{\sigma_\lambda^2}{\lambda_0^2}\\
&=\left(-\frac{x^2}{4}\frac{(1-x)\ln\left(\frac{2}{1-x}\right)^2+(1+x)\ln\left(\frac{2}{1+x}\right)^2}{(1-x)\ln\left(\frac{2}{1-x}\right)+(1+x)\ln\left(\frac{2}{1+x}\right)}-\frac{(1-x)x}{8}\ln\left(\frac{2}{1-x}\right)+\frac{(1+x)x}{8}\ln\left(\frac{2}{1+x}\right)\right)\frac{\sigma_\lambda^2}{\lambda_0^2},
\end{aligned}\tag{41}$$

$$A_1=\left(-x^2\frac{(1-x)\ln\left(\frac{2}{1-x}\right)^2+(1+x)\ln\left(\frac{2}{1+x}\right)^2}{\left((1-x)\ln\left(\frac{2}{1-x}\right)+(1+x)\ln\left(\frac{2}{1+x}\right)\right)^2}+\frac{x}{2}\frac{-(1-x)\ln\left(\frac{2}{1-x}\right)+(1+x)\ln\left(\frac{2}{1+x}\right)}{(1-x)\ln\left(\frac{2}{1-x}\right)+(1+x)\ln\left(\frac{2}{1+x}\right)}\right)\frac{\sigma_\lambda^2}{\lambda_0^2}.\tag{42}$$

Therefore,

$$\begin{aligned}
\Lambda_p=\lambda_0&\left[1+\left(-1+\frac{1}{2}\frac{(1-x)\ln\left(\frac{2}{1-x}\right)^2+(1+x)\ln\left(\frac{2}{1+x}\right)^2}{(1-x)\ln\left(\frac{2}{1-x}\right)+(1+x)\ln\left(\frac{2}{1+x}\right)}\right)\frac{\sigma_\lambda^2}{\lambda_0^2}\right.\\
&\left.+(1-\alpha)\left(-x^2\frac{(1-x)\ln\left(\frac{2}{1-x}\right)^2+(1+x)\ln\left(\frac{2}{1+x}\right)^2}{\left((1-x)\ln\left(\frac{2}{1-x}\right)+(1+x)\ln\left(\frac{2}{1+x}\right)\right)^2}+\frac{x}{2}\frac{-(1-x)\ln\left(\frac{2}{1-x}\right)+(1+x)\ln\left(\frac{2}{1+x}\right)}{(1-x)\ln\left(\frac{2}{1-x}\right)+(1+x)\ln\left(\frac{2}{1+x}\right)}\right)\frac{\sigma_\lambda^2}{\lambda_0^2}\right]+O(\delta\alpha^2).
\end{aligned}\tag{43}$$

### 5 Tunable Inhibitor Dilution Model

Here we discuss the inhibitor dilution model applied in Figure S1 (C). In this model, cell cycle progression is limited by the dilution of an inhibitor molecule to a critical concentration. Within this modeling framework, we consider a cell cycle that is split into two phases: the G1 phase, and the budded phase. Within this model, cells direct new growth in the budded phase to a newly forming bud that will separate to form a new $D$ type cell at mitosis, leaving the main cell as the $M$ type cell. For simplicity, we consider the duration of the budded phase to be constant, generating an approximately constant division asymmetry $x$ between $M$ and $D$ cells. During the budded phase, cells also produce an amount $\Delta$ of some inhibitor $I$. In the G1 phase of the subsequent cell cycle, this inhibitor is then diluted through growth until it reaches a critical concentration, causing the cell to initiate the budded phase. We have considered this biologically relevant model in greater depth elsewhere [5, 6]. Here we consider a broader class of models in which a cell degrades some fraction $a$ of its inhibitor immediately after passing through Start. These definitions lead to the relations

$$\begin{aligned} I_d &= (1-a)I_b + \Delta, \\ V_d^D &= (1-a)V_b^D + \Delta\frac{1-x}{1+x}, \\ V_d^M &= (1-a)V_b^M + \Delta. \end{aligned} \tag{44}$$

Varying $a$ therefore tunes the size regulation behavior from an adder for $a = 0$ to a sizer for $a = 1$. Motivated by the previous results, we tested the behavior of $\Lambda_P$ for variable $a$, $\sigma_\lambda$, and $x$. We also tested the addition of time additive noise $\sigma_t$ in passage through start. Figure S1 (C) demonstrates that the results for the sizer model from Section S3 carry over to the $a = 1$ case, with noise in $\lambda$ decreasing the population growth rate, and greater division asymmetry offsetting this deficit.

### 6 Testing the Euler-Lotka equation for non-IGT growth policies

Here we evaluate the accuracy of the Euler-Lotka equation (Equation 1 in the main text) for non-IGT growth policies using numerical estimations. We performed simulations across a range of size control strategies by varying $\alpha$ in Equation 4 in the main text, as well as the division asymmetry $x$, and calculated the population growth rate directly in each case. Additionally, we calculated a kernel density estimate (KDE) of the generation time distribution for each simulated population using the Python sklearn package. By inserting this KDE into Equation S10 and computing the corresponding integral we inferred the corresponding value of $\Lambda_p$ for that KDE. The results of this comparison are summarized in Figure S5, and demonstrate strong agreement between the simulated value of $\Lambda_p$ and the numerically estimated value across all tested parameter sets.

### 7 Growth rate penalty model

Here we consider deviations from I.I.D. growth rates. We assume a noisy single cell growth rate $\lambda$, the average of which is dependent on cell volume at birth,

$$\lambda \sim \mathcal{N}\left(\lambda_0 f\left(\frac{V_b - V^*}{V^*}\right), \sigma_\lambda^2\right) \tag{45}$$

for a given volume at birth $V_b$, where $f((V_b - V^*)/V^*)$ is some function with a maximum at $V_b = V^*$, so that $V^*$ represents the "optimal" cell size at birth with the largest average single cell growth rate. This model directly couples the distribution of growth rates measured throughout the population to the distribution of volumes at birth. We will adopt the simplifying assumption that $f$ is symmetric about $V_b = V^*$, and that $V^* = \langle V_b \rangle_P$ is the population average cell volume at birth, calculated over the full population tree. As in Section S3 we can make progress analytically on this model in the case of a sizer with $\sigma_t = 0$. In this case, the growth rate simplifies to an identical expression for all cells:

$$\lambda \sim N\left(\lambda_0 f(x), \sigma_\lambda^2\right), \tag{46}$$

with the average growth rate decreasing with increasing division asymmetry $x$. This change amounts to the substitution of $\lambda_0 \rightarrow \lambda_0 f(x)$ in Equation S18.

As an example, we consider the case $f(x) = 1 - \epsilon x^n$, with penalty strength $\epsilon$ and exponent $n$. The case $n = 0$ corresponds to a constant average growth rate independent of cell volume, and is identical to the model studied in Section 3 under the appropriate normalization $\lambda_0 \to \lambda_0 - \epsilon$. The restrictions that $f(x)$ is symmetric and has a maximum at $x = 0$ lead us to focus on the behavior arising when $n = 2$ or $n = 4$, with

$$\begin{aligned}\Lambda_P(\sigma_\lambda, x) \approx &\lambda_0(1 - \epsilon x^n) \times \\ &\left(1 - \left(1 + \frac{1}{2}\frac{(1+x)(\ln(\frac{1+x}{2}))^2 + (1-x)(\ln(\frac{1-x}{2}))^2}{(1+x)\ln(\frac{1+x}{2}) + (1-x)\ln(\frac{1-x}{2})}\right)\left(\frac{\sigma_\lambda}{\lambda_0(1-\epsilon x^n)}\right)^2\right)\end{aligned} \tag{47}$$

Figure S6 shows the agreement of Equation S47 with simulations for (A) the $n = 2$ case, and (B) the $n = 4$ case, with increasing penalty strength $\epsilon$. Increasing $\epsilon$ for $n = 2$ causes $\Lambda_P$ to decrease initially before rising again for large $x$, creating a local maximum in $\Lambda_P$ at $x = 0$. In contrast, for $n = 4$ we observe a local maximum in $\Lambda_P$ at nonzero $x$. It is to be expected that our introduction of a growth rate penalty associated with broad cell size distributions should generate local maxima in $\Lambda_P$ at $x = 0$, however, it is notable that the generation of local maxima at finite $x$ only occurs if the $n = 4$ coefficient is nonzero.

We used simulations to study the behavior of $\Lambda_P$ for non-sizer growth policies. Figure S6 demonstrates the behavior of non-sizer growth policies upon varying division asymmetry $x$ for a fixed level of growth rate noise $\sigma_\lambda/\lambda_0 = 0.1$ and $\sigma_t = 0$. Weaker size control strategies show a pronounced decrease in growth rate for large $x$ as shown. This decrease in $\Lambda_P$ arises from a broadening of the cell size distribution for such weaker strategies, which consequently incurs a stronger penalty from the decrease in growth rate for excessively large or small cells in Equation S45.

### 8 Growth rate correlations

Here we present a model for correlated growth rates across generations. Following [7], we assume that the single cell exponential growth rate in the $n+1^{th}$ generation, $\lambda_{n+1}$, follows

$$\lambda_{n+1} = a\lambda_n + b + \eta, \tag{48}$$

where $\eta \sim N(0, \sigma_\epsilon)$ is a random, I.I.D. noise. The Pearson correlation coefficient is then given by the parameter $a \in (0, 1)$. The average growth rate is then $\langle\lambda\rangle = b/(1-a)$, while the variance is given by $\sigma_\lambda^2 = \sigma_\epsilon^2/(1-a^2)$. Results for simulations of this model are plotted in Figure S7.

### 9 Positive generation time correlations

Here we present the result that size control in asymmetrically dividing cells can lead to positive generation time correlations between parents and their progeny. It has been shown previously in symmetrically dividing cells that at steady state the Pearson correlation coefficient (PCC) between the generation times of a parent cell and its progeny that is predicted by Equation 7 in the main text is negative, with $PCC \equiv \frac{\langle \tau_{n+1}\tau_n\rangle_c}{\sigma_\tau \sigma_{\tau_{n+1}}} = -\frac{\alpha}{2}$ [1]. This negative correlation is intuitive, since a cell with an unusually long division time in one generation will produce larger progeny that will compensate with a correspondingly a shorter division time.

In the next section we derive an analytical expression for the PCC of asymmetrically dividing cells based on Equation 7 in the main text. For the PCC between the generation time of a daughter or mother cell and the generation time of its parent in the previous cell cycle this takes the form

$$PCC(n, n+1) = \frac{\alpha\left((1-\alpha)\left(\frac{\ln\left(\frac{1-x}{1+x}\right)}{2\lambda}\right)^2 - \sigma_t^2\right)}{\sqrt{\left(2\sigma_t^2 + \alpha\left(\frac{\ln\left(\frac{1-x}{1+x}\right)}{2\lambda}\right)^2\right)\left(2\sigma_t^2 + \alpha(1-\alpha)^2\left(\frac{\ln\left(\frac{1-x}{1+x}\right)}{2\lambda}\right)^2\right)}}. \tag{49}$$

The expression is identical for mother or daughter cells. For symmetric division ($x = 0$) we have the further simplification that $PCC = -\frac{\alpha}{2}$, as derived in [1]. The PCC between the generation time of the two cells (one daughter and one mother) generated by a single cell division event is given by

$$PCC(n_M, n_D) = \frac{\langle \tau_{n+1,D}\tau_{n+1,M}\rangle_c}{\sigma^2_{\tau_{n+1}}} = 1 - \frac{(2-\alpha)\sigma_t^2}{2\sigma_t^2 + \alpha(1-\alpha)^2\left(\frac{\ln\left(\frac{1-x}{1+x}\right)}{2\lambda}\right)^2}. \tag{50}$$

Figure S4 shows good agreement between these predictions and our simulations. We note that the agreement becomes poor for high $\sigma_t$ and $x$ due to a small fraction of mother cells becoming sufficiently large for their generation times to be extremely small in the absence of noise. In this case the assumption that $\eta$ is a Gaussian noise breaks down, since generation time must always be positive. We note that for simplicity we assumed $\sigma_\lambda = 0$ in the derivation of Equations S49 and S50. As such, we used simulations to explore the effect of noise in $\lambda$ on these correlations, with results plotted in Figure S4 (C, F). Increasing $\sigma_\lambda$ decreases the strength of the correlations, but for biologically relevant noise strengths with $\sigma_\lambda/\lambda_0 \approx 0.1-0.2$, asymmetric division can still generate positive correlations in both $PCC(n, n+1)$ and $PCC(n_M, n_D)$.

#### 9.1 Calculations of Generation Time Correlations

The Pearson correlation coefficient between the generation time of a parent cell and its progeny (either mother or daughter) is given by

$$PCC = \frac{\langle \tau_{n+1}\tau_n\rangle_c}{\sigma_\tau \sigma_{\tau_{n+1}}} \tag{51}$$

Here $\sigma_\tau$ corresponds to the standard deviation in generation times over both cell types, while $\sigma_{\tau_{n+1}}$ is the standard deviation for either the daughter or mother cell types (the expression is identical for each type). We note that $\langle XY\rangle_c \equiv \langle (X - \langle X\rangle)(Y - \langle Y\rangle)\rangle$.

We now calculate $\langle \tau_{n+1}\tau_n\rangle_c \equiv \langle \tau_{n+1}\tau_n\rangle - \langle \tau_{n+1}\rangle\langle \tau_n\rangle$ for either a mother or daughter cell in the $n+1^{th}$ generation, born from a randomly sampled cell of unspecified cell type in the $n^{th}$ generation. We assume a constant strength of time-additive noise $\sigma_t^2$ is kept constant for both mothers and daughters. Applying Equation 7 from the main text yields

$$\begin{aligned}
\tau_{n+1} &= \frac{\ln 2}{\lambda} - \frac{\alpha}{\lambda}\ln|\frac{v_{b,n+1}}{\Delta}| + \eta' \\
&= \frac{\ln 2}{\lambda} - \frac{\alpha}{\lambda}\ln|\frac{(1-x)v_{b,n}e^{\lambda\tau_n}}{2\Delta}| + \eta' \\
&= \frac{\ln 2}{\lambda} - \frac{\alpha}{\lambda}(\ln(\frac{1-x}{2}) + \ln|\frac{v_{b,n}}{\Delta}|) - \alpha\tau_n + \eta' \\
\Rightarrow \langle \tau_{n+1}\tau_n\rangle_c &= \frac{\alpha^2}{\lambda^2}\langle \ln|\frac{v_{b,n}}{\Delta}|^2\rangle_c - \alpha\langle \tau_n^2\rangle_c.
\end{aligned} \tag{52}$$

Note that this calculation is carried out here for a $D$ cell in the $n+1^{th}$ generation, but the result also holds for an $M$ type cell since the result is independent of $x$. We take $\ln|\frac{v_{b,n}}{\Delta}| \equiv \omega$, so that

$$\langle \tau_{n+1}\tau_n \rangle_c = \frac{\alpha^2}{\lambda^2}\sigma_\omega^2 - \alpha\sigma_\tau^2. \tag{53}$$

We apply Equation 7 again to obtain the generation time variance

$$\begin{aligned}\sigma_\tau^2 &= (\frac{\ln 2}{\lambda})^2 + \frac{\alpha^2}{\lambda^2}\langle\omega^2\rangle + \sigma_t^2 - \frac{2\alpha}{\lambda^2}\langle\omega\rangle \ln 2 - (\frac{\ln 2}{\lambda} - \frac{\alpha}{\lambda}\langle\omega\rangle)^2 \\ &= \frac{\alpha^2}{\lambda^2}\sigma_\omega^2 + \sigma_t^2.\end{aligned} \tag{54}$$

Here $\sigma_t$ is the standard deviation of time additive noise. We now calculate

$$\begin{aligned}\omega_D^{n+1} &= \ln\left|\frac{(1-x)v_{b,n}e^{\lambda\tau_n}}{2\Delta}\right| \\ \Rightarrow \langle\omega_D\rangle &= \ln|(1-x)| + (1-\alpha)\langle\omega\rangle \\ \Rightarrow \langle\omega_M\rangle &= \ln|(1+x)| + (1-\alpha)\langle\omega\rangle.\end{aligned} \tag{55}$$

The last line is obtained by substituting $1-x \to 1+x$ to interchange results from daughter cells to mother cells. We therefore obtain

$$\begin{aligned}\langle\omega\rangle &= \frac{1}{2}(\langle\omega'_D\rangle + \langle\omega'_M\rangle) \\ &= \frac{1}{2\alpha}\ln|(1-x)(1+x)|.\end{aligned} \tag{56}$$

Similarly,

$$\begin{aligned}\langle\omega_D^2\rangle &= \ln|1-x|^2 + 2(1-\alpha)\ln|1-x|\langle\omega\rangle + (1-\alpha)^2\langle\omega^2\rangle + \lambda^2\sigma_t^2 \\ \langle\omega_M^2\rangle &= \ln|1+x|^2 + 2(1-\alpha)\ln|1+x|\langle\omega\rangle + (1-\alpha)^2\langle\omega^2\rangle + \lambda^2\sigma_t^2 \\ \langle\omega^2\rangle &= \frac{1}{2}(\langle\omega_D^2\rangle + \langle\omega_M^2\rangle) \\ \Rightarrow \alpha(2-\alpha)\langle\omega^2\rangle &= \frac{1}{2}(\ln|1-x|^2 + \ln|1+x|^2) + \frac{(1-\alpha)}{2\alpha}\ln|(1-x)(1+x)|^2 + \lambda^2\sigma_t^2 \\ \Rightarrow \langle\omega^2\rangle &= \frac{1}{2\alpha(2-\alpha)}(\ln|1-x|^2 + \ln|1+x|^2) + \frac{(1-\alpha)}{2\alpha^2(2-\alpha)}\ln|(1-x)(1+x)|^2 \\ &\quad + \frac{\lambda^2\sigma_t^2}{\alpha(2-\alpha)} \\ \Rightarrow \sigma_\omega^2 &= \langle\omega^2\rangle - \langle\omega\rangle^2 \\ &= -\frac{1}{4\alpha(2-\alpha)}\ln|(1-x)(1+x)|^2 + \frac{1}{2\alpha(2-\alpha)}(\ln|1-x|^2 + \ln|1+x|^2) \\ &\quad + \frac{\lambda^2\sigma_t^2}{\alpha(2-\alpha)} \\ &= \frac{1}{4\alpha(2-\alpha)}\ln|\frac{1-x}{1+x}|^2 + \frac{\lambda^2\sigma_t^2}{\alpha(2-\alpha)} \\ \Rightarrow \sigma_\omega^2 &= \frac{1}{\alpha(2-\alpha)}\left(\frac{1}{4}\ln|\frac{1-x}{1+x}|^2 + \lambda^2\sigma_t^2\right)\end{aligned} \tag{57}$$

Note that $x=0$ for symmetric division yields $\sigma_\omega^2 = \lambda^2\sigma_t^2/(\alpha(2-\alpha))$ as in [8]. We can now insert this form into Equation S54 to find

$$\sigma_\tau^2 = \frac{2\sigma_t^2}{2-\alpha} + \frac{\alpha}{2-\alpha}\left(\frac{\ln\frac{1-x}{1+x}}{2\lambda}\right)^2. \tag{58}$$

We now can substitute into Equation S53 to obtain

$$\langle \tau_{n+1}\tau_n\rangle_c = \frac{\alpha}{2-\alpha}\left((1-\alpha)\left(\frac{\ln\frac{1-x}{1+x}}{2\lambda}\right)^2 - \sigma_t^2\right) \tag{59}$$

This value is positive for $\sigma_t^2 < (1-\alpha)(\ln\frac{1-x}{1+x}/(2\lambda))^2$, with $\alpha < 1$, contrasting with previously observed results in symmetric division.

To calculate $\sigma^2_{\tau_{n+1}}$ we take

$$\begin{aligned}
\sigma^2_{\tau_{n+1}} &= \left\langle \left(\frac{\ln 2}{\lambda} - \frac{\alpha(\ln\left|\frac{1-x}{2}\right| + \omega)}{\lambda} - \alpha\tau_n + \eta'\right)^2 \right\rangle_c \\
&= \frac{\alpha^2\sigma_\omega^2}{\lambda^2} + \frac{2\alpha^2\langle\omega_n\tau_n\rangle_c}{\lambda} + \alpha^2\sigma_\tau^2 + \sigma_t^2 \\
&= \frac{\alpha^2\sigma_\omega^2(1-2\alpha)}{\lambda^2} + \alpha^2\sigma_\tau^2 + \sigma_t^2 \\
&= \frac{2\sigma_t^2 + \alpha(1-\alpha)^2\left(\frac{\ln\frac{1-x}{1+x}}{2\lambda}\right)^2}{2-\alpha}.
\end{aligned} \tag{60}$$

The third line follows since $\langle\omega_n\tau_n\rangle_c = -\frac{\alpha}{\lambda}\sigma_\omega^2$, while the final equality follows from substituting in the expressions for $\sigma_\omega^2$ and $\sigma_\tau$. Note that this expression is not the same as that in Equation S58 because we have conditioned it on the cell in the $n+1^{th}$ generation being a specific cell type rather than being drawn from the full population distribution, even though the result is independent of whether that specific cell type is a daughter or mother.

For comparison with [3] we also evaluate the correlation between the generation times of the two progeny from any given cell division event.

To calculate $\langle\tau_{n+1,D}\tau_{n+1,M}\rangle_c$ we take

$$\begin{aligned}
\langle\tau_{n+1,D}\tau_{n+1,M}\rangle_c &= \langle\frac{\alpha^2\omega_n^2}{\lambda^2}\rangle_c + \alpha^2\sigma_\tau^2 + \frac{2\alpha^2}{\lambda}\langle\omega_n\tau_n\rangle_c \\
&= \alpha^2\sigma_\tau^2 + \frac{\alpha^2\sigma_\omega^2}{\lambda^2}(1-2\alpha) \\
&= \sigma^2_{\tau_{n+1}} - \sigma_t^2 \\
&= \frac{\alpha\sigma_t^2 + \alpha(1-\alpha)^2\left(\frac{\ln\frac{1-x}{1+x}}{2\lambda}\right)^2}{2-\alpha}
\end{aligned} \tag{61}$$

We therefore have

$$PCC(mother-daughter) = \frac{\langle\tau_{n+1,D}\tau_{n+1,M}\rangle_c}{\sigma^2_{\tau_{n+1}}} = 1 - \frac{\sigma_t^2}{\sigma^2_{\tau_{n+1}}}, \tag{62}$$

or equivalently

$$PCC = \frac{\langle\tau_{n+1,D}\tau_{n+1,M}\rangle_c}{\sigma^2_{\tau_{n+1}}} = 1 - \frac{(2-\alpha)\sigma_t^2}{2\sigma_t^2 + \alpha(1-\alpha)^2\left(\frac{\ln\left(\frac{1-x}{1+x}\right)}{2\lambda}\right)^2}. \tag{63}$$

### 10 Supplementary Figures

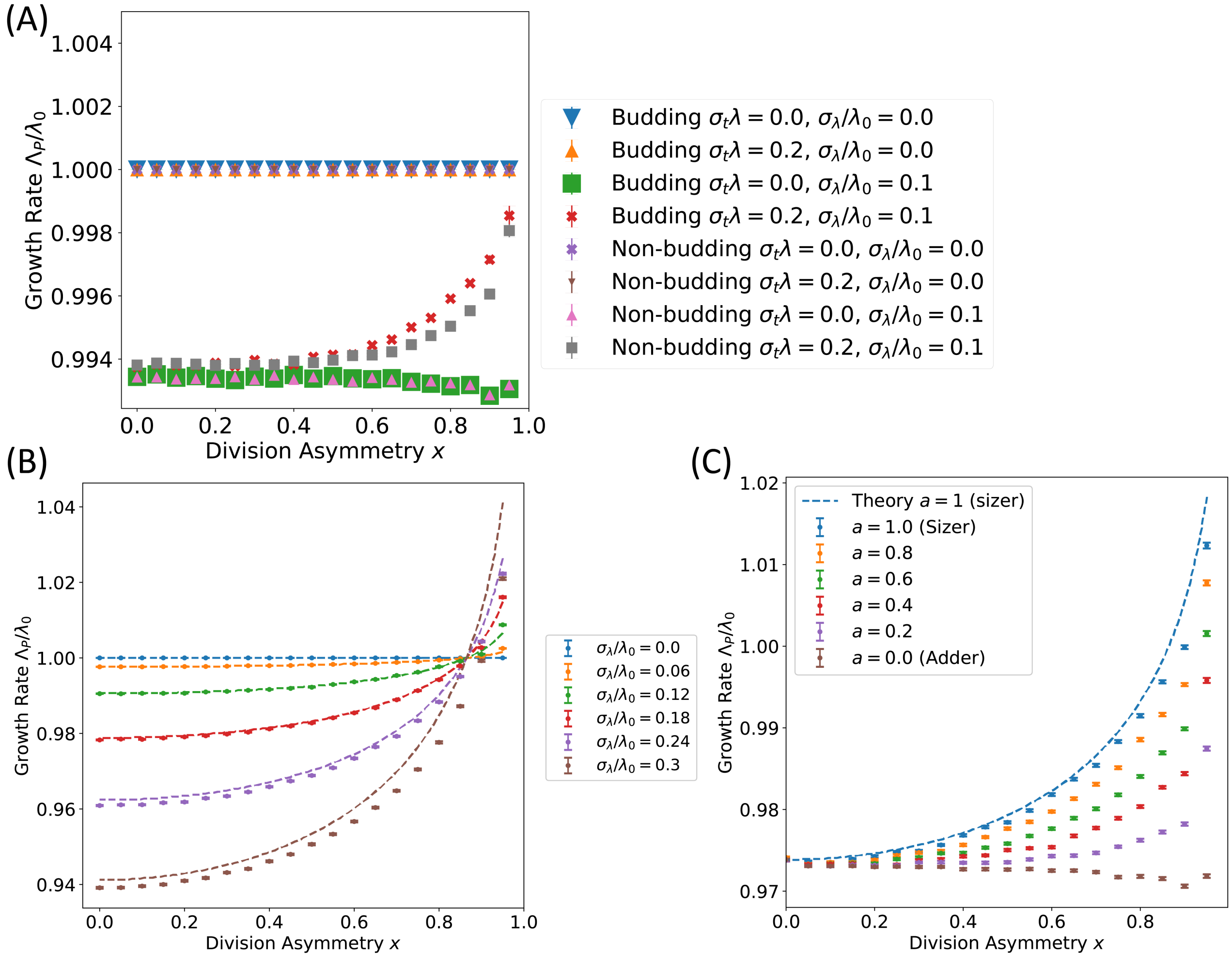


Figure 1: (A) Simulated values for $\Lambda_P$ plotted against division asymmetry $x$ for an adder model ($\alpha = 0.5$) following either budding or non-budding growth morphologies. Parameters match those of Figure 2 (A) in the main text. Generation time noise causes a substantial effect on the population growth rate in the regime of extreme division asymmetries, and we observe minor deviations between budding and non-budding growth morphologies in this regime. (B) Comparison of Equation 5 in the main text with simulations for a sizer model ($\alpha = 1$) shows good agreement for small $\sigma_\lambda$. (C) $\Lambda_P$ for cells simulated according to a tunable inhibitor dilution model discussed in Section S5. $\Lambda_P$ shows similar dependency on $x$, $\sigma_\lambda$ and size control strategy to that shown in Figure 2 (B) in the main text. Plotted for $\sigma_\lambda/\lambda_0 = 0.2$, $\sigma_t = 0$.

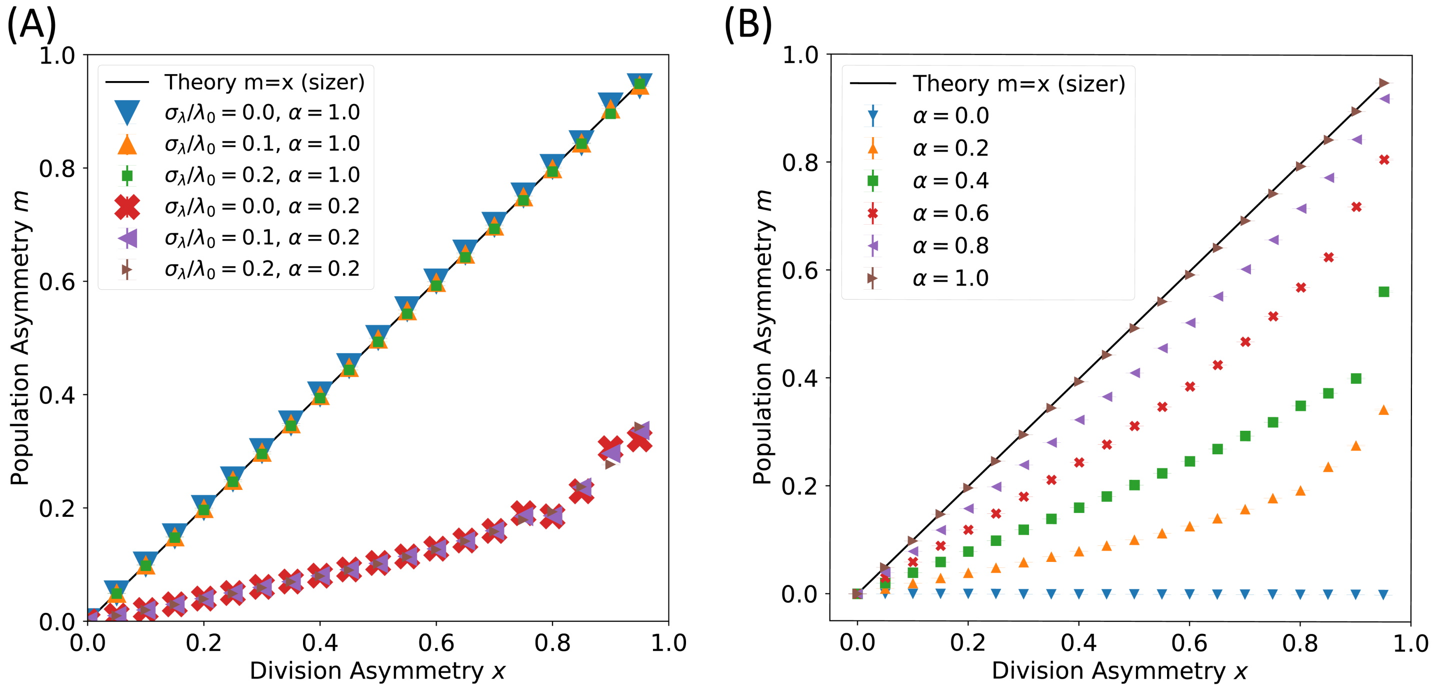


Figure 2: The asymmetry $m = (N_D - N_M)/(N_D + N_M)$ in population size between mothers and daughters at a given point in time increases with increasing division asymmetry in a manner that depends on the strategy of size control, and is independent of noise in the growth rate $\sigma_\lambda$. (A) $m$ plotted against asymmetry $x$ for a sizer $\alpha = 1$ agrees with $m = x$ for a range of $\sigma_\lambda$ values. $m$ plotted for $\alpha = 0.2$ shows that $m$ is independent of $\sigma_\lambda$ for all size control strategies tested. (B) $m$ plotted against asymmetry $x$ for a range of size control strategies $\alpha$ show that weaker size control strategies have a decreased value of $m$.

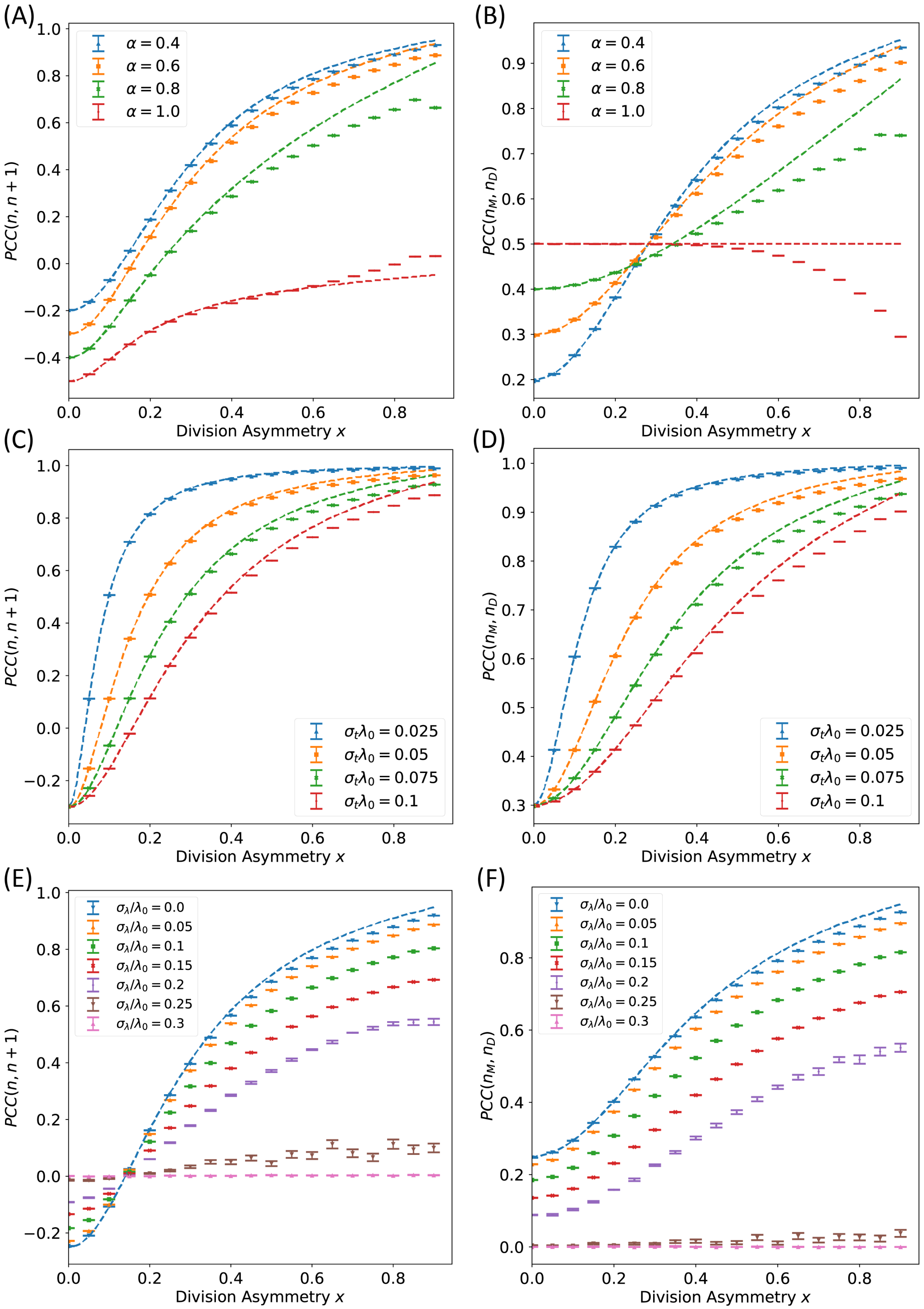


Figure 3: Figure legend continued on next page.

Figure 3: Asymmetric division generates positive correlations between closeley related cells. (A,C,E) Correlation coefficients for the generation times of daughter cells and those of their parent cells, $PCC(n, n+1)$, plotted against division asymmetry $x$ for a range of size control strategies and noise strengths. (B,D,F) Correlation coefficients for the generation times of the daughter and mother cells generated by a cell division event ($PCC(n_M, n_D)$) plotted against division asymmetry $x$ for a range of size control strategies and noise strengths. (A) PCC plotted for variable size control strategy $\alpha$ as shown, with $\sigma_\lambda = 0$ and $\sigma_t\lambda_0 = 0.1$. (C) PCC plotted for variable generation time noise $\sigma_t$ as shown, with $\sigma_\lambda = 0$ and $\alpha = 0.6$. (E) PCC plotted for variable $\sigma_\lambda$ as shown, with $\sigma_t\lambda_0 = 0.1$ and $\alpha = 0.5$. (B) PCC plotted for variable size control strategy $\alpha$ as shown, with $\sigma_\lambda = 0$ and $\sigma_t\lambda_0 = 0.1$. (D) PCC plotted for variable generation time noise $\sigma_t$ as shown, with $\sigma_\lambda = 0$ and $\alpha = 0.6$. (F) PCC plotted for variable $\sigma_\lambda$ as shown, with $\sigma_t\lambda_0 = 0.1$ and $\alpha = 1/2$. Data points correspond to simulations, while dotted lines represent theory predictions. Error bars show the standard error of the mean over 100 simulated repeats.

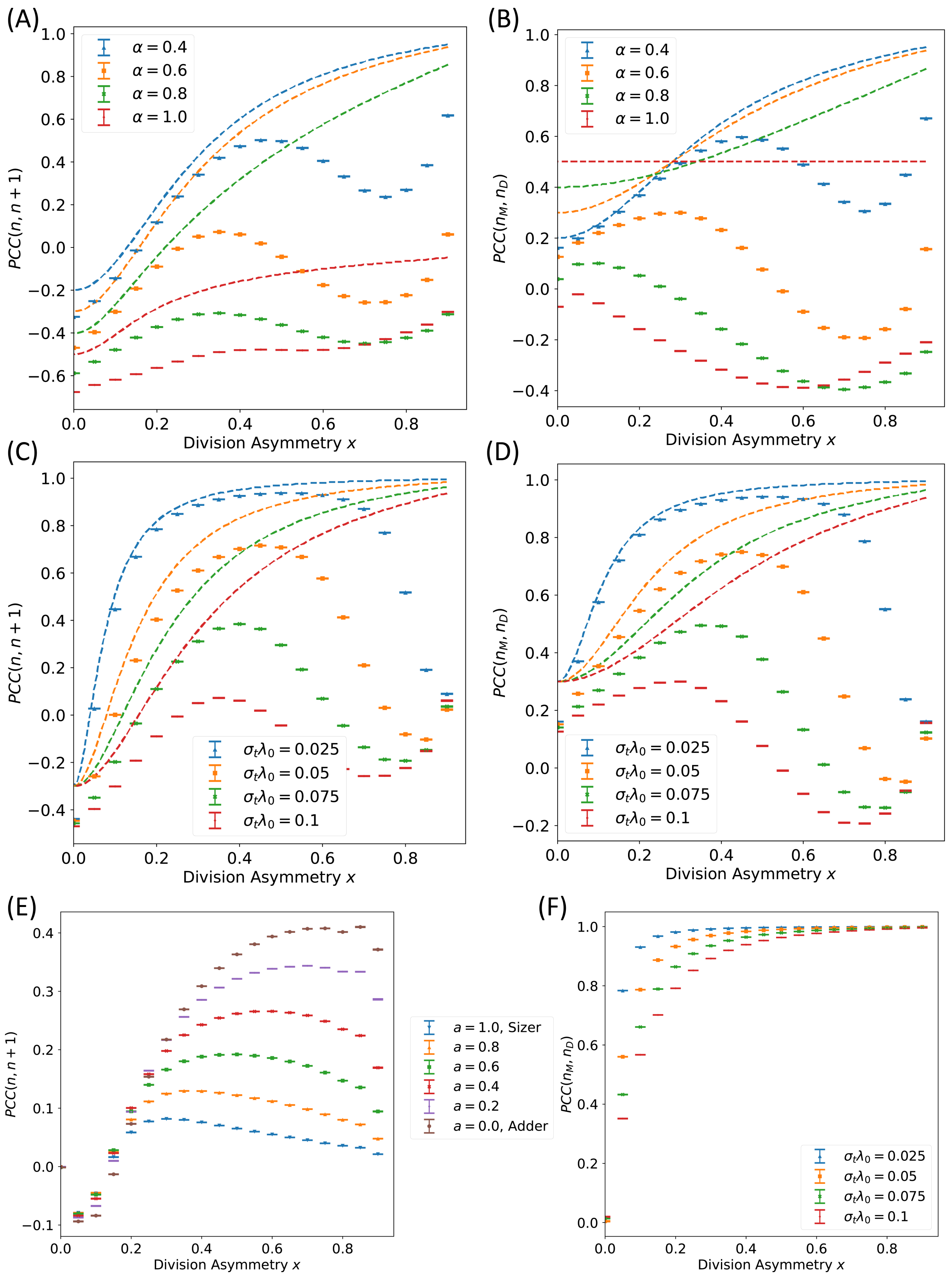


Figure 4: Figure legend continued on next page.

Figure 4: Cells that have a budding growth morphology maintain positive generation time correlations in a broad regime of parameter space, while showing significant deviations from the correlations of non-budding cells for increasing $\sigma_t$, $\alpha$ and $x$ (cf. Fig. S3). (A,C) Correlation coefficients for the generation times of daughter cells and those of their parent cells, $PCC(n, n+1)$, plotted against division asymmetry $x$ for a range of size control strategies and noise strengths. (B,D) Correlation coefficients for the generation times of the daughter and mother cells generated by a cell division event ($PCC(n_M, n_D)$) plotted against division asymmetry $x$ for a range of size control strategies and noise strengths. (A) PCC plotted for variable size control strategy $\alpha$ as shown, with $\sigma_\lambda = 0$ and $\sigma_t\lambda_0 = 0.1$. (C) PCC plotted for variable generation time noise $\sigma_t$ as shown, with $\sigma_\lambda = 0$ and $\alpha = 0.6$. (B) PCC plotted for variable size control strategy $\alpha$ as shown, with $\sigma_\lambda = 0$ and $\sigma_t\lambda_0 = 0.1$. (D) PCC plotted for variable generation time noise $\sigma_t$ as shown, with $\sigma_\lambda = 0$ and $\alpha = 0.6$. Data points correspond to simulations, while dotted lines represent theory predictions. Error bars show the standard error of the mean over 100 simulated repeats. Cells were simulated to grow with a budding growth morphology. (E-F) Correlation coefficients for the generation times of cells simulated using an inhibitor dilution model for cell cycle progression, and dividing with a non-budding morphology. (E) Correlation coefficients for the generation times of daughter cells and those of their parent cells, $PCC(n, n+1)$, plotted against division asymmetry $x$ for a range of size control strategies ranging between a sizer for $a = 1.0$ and an adder for $a = 0.0$. $\sigma_\lambda = 0$, and $\sigma_t\lambda_0 = 0.1$. (F) Correlation coefficients for the generation times of daughter and mother cells generated by a cell division event ($PCC(n_M, n_D)$) plotted against division asymmetry $x$ for a range of size control strategies ranging between a sizer for $a = 1.0$ and an adder for $a = 0.0$. $\sigma_\lambda = 0$, and $\sigma_t\lambda_0 = 0.1$.

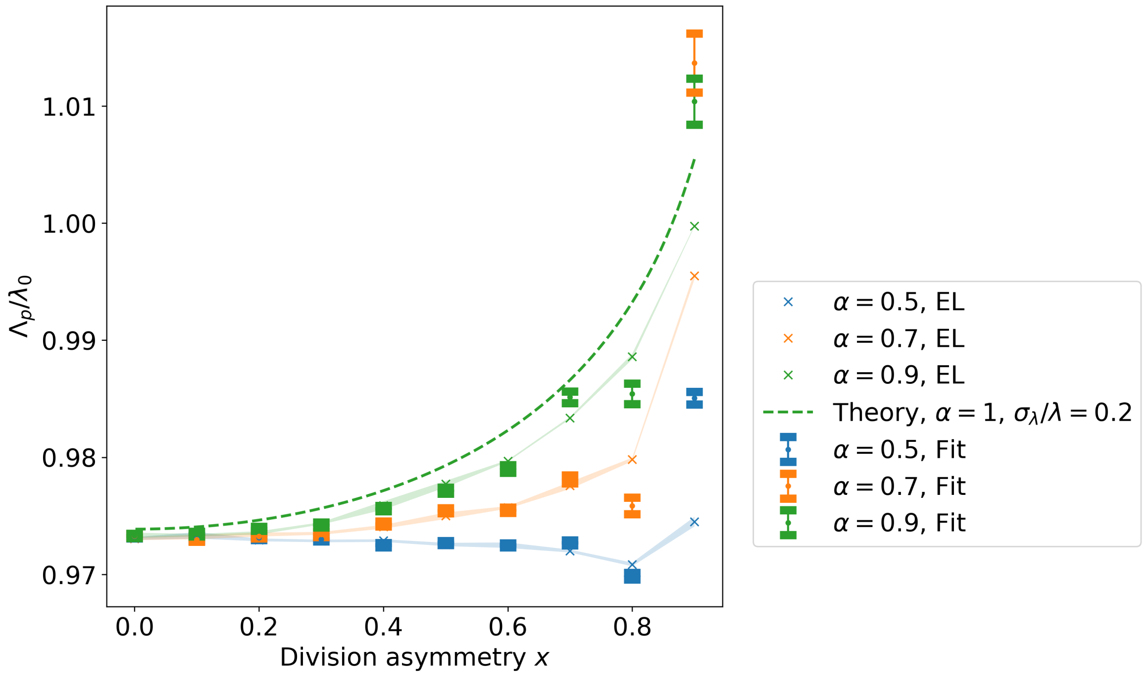


Figure 5: Comparison of simulated values for $\Lambda_p$ vs. numerically estimated values using Equation S10 across a range of parameter values, as described in Section S6. Simulated growth rates based on a fit to the exponential growth of the cell population are plotted with error bars, while the growth rate calculated from numerical integration of the Euler-Lotka equation is shown as a coloured line. Error bars show standard error of the mean. The simulations show strong agreement with the prediction of Equation 10 throughout. Results were obtained with $\sigma_\lambda/\lambda_0 = 0.2$, $\sigma_t = 0$.

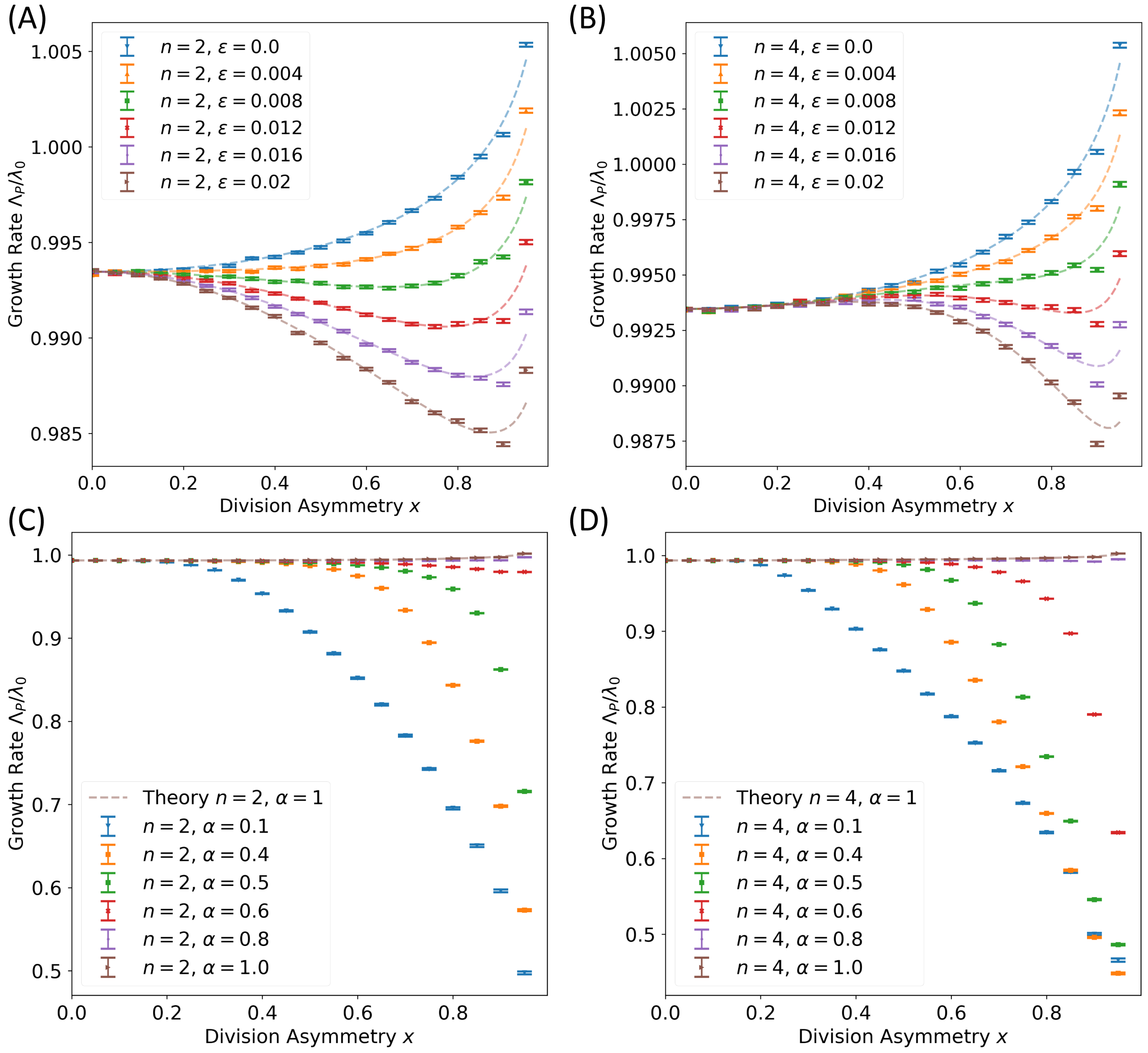


Figure 6: Plots of the population growth rate $\Lambda_P$ for cells undergoing asymmetric division and growing with a growth rate penalty described by Equation S45, given $f(x) = 1 - \epsilon x^n$. Introducing a growth rate penalty generates local maxima in $\Lambda_P$ at $x = 0$ for $n = 2$, and at finite $x$ for $n = 4$. (A-B) Predicted $\Lambda_P$ for cells growing according to a Sizer size control strategy with varying penalty strength $\epsilon$ show good agreement with the predictions of Equation S47. Simulations are generated for $\sigma_\lambda/\lambda_0 = 0.1$ and $\sigma_t = 0.0$. (C-D) Simulated predictions for cells growing with variable size control strategies for $\epsilon = 0.04$, $\sigma_\lambda/\lambda_0 = 0.1$ and $\sigma_t = 0.0$. The growth rate penalty becomes more pronounced for asymmetrically dividing cells with weaker size control strategies. Results are plotted for variable $x$, with (A, C) $n = 2$ and (B, D) $n = 4$. Data points correspond to simulations, while dotted lines represent theory predictions. Error bars show the standard error of the mean over 100 simulated repeats.

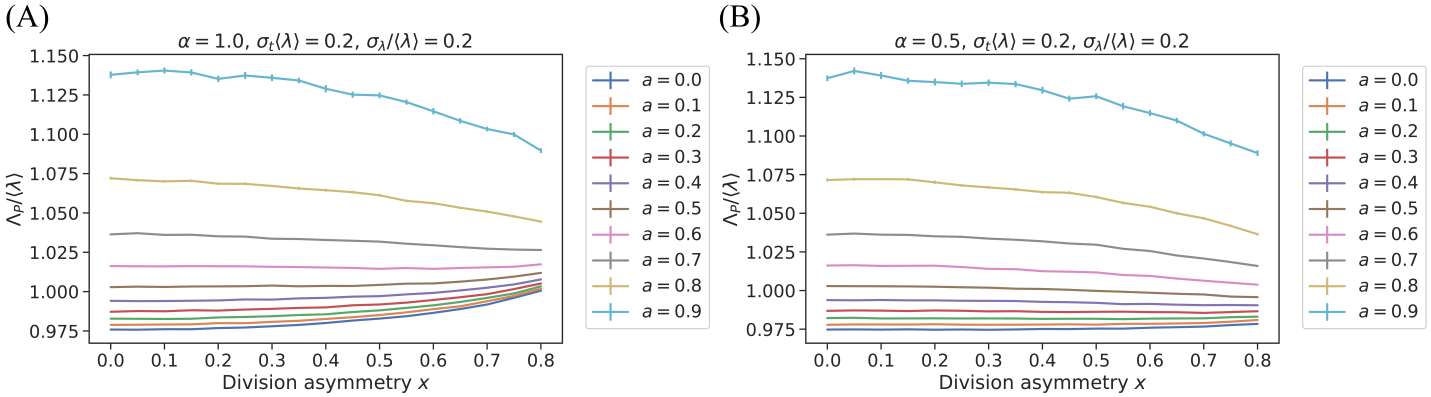


Figure 7: Plots of the population growth rate $\Lambda_P$ for cells with growth rate correlations implemented according to Equation S48. Growth rate correlations have strength determined by the parameter $a$. Plots show results for cells simulated to follow (A) a sizer size control strategy, or (B) an adder size control strategy. Error bars show the standard error of the mean over 100 simulated repeats.

[6] Felix Barber, Po-Yi Ho, Andrew W. Murray, and Ariel Amir. Details Matter: Noise and Model Structure Set the Relationship between Cell Size and Cell Cycle Timing. *Frontiers in Cell and Developmental Biology*, 5, 2017.

[7] Jie Lin and Ariel Amir. From single-cell variability to population growth. *Physical Review E*, 101(1):012401, 2020.

[8] Ariel Amir. Cell Size Regulation in Bacteria. *Physical Review Letters*, 112(20):208102, 2014.
